# Increased excitability of dentate gyrus mossy cells occurs early in life in the Tg2576 model of Alzheimer’s disease

**DOI:** 10.1101/2024.02.09.579729

**Authors:** David Alcantara-Gonzalez, Meghan Kennedy, Chiara Criscuolo, Justin Botterill, Helen E Scharfman

**Author notes:** Corresponding authors: David Alcantara-Gonzalez Helen Scharfman Center for Dementia Research 140 Old Orangeburg Rd. Bldg. 39 Orangeburg, NY 10962.

## Abstract

**INTRODUCTION:** Hyperexcitability in Alzheimer’s disease (AD) emerge early and contribute to disease progression. The dentate gyrus (DG) is implicated in hyperexcitability in AD. We hypothesized that mossy cells (MCs), regulators of DG excitability, contribute to early hyperexcitability in AD. Indeed, MCs generate hyperexcitability in epilepsy.

**METHODS:** Using the Tg2576 model and WT mice (∼1month-old), we compared MCs electrophysiologically, assessed c-Fos activity marker, Aβ expression and mice performance in a hippocampal-dependent memory task.

**RESULTS:** Tg2576 MCs exhibit increased spontaneous excitatory events and decreased inhibitory currents, increasing the charge transfer excitation/inhibition ratio. Tg2576 MC intrinsic excitability was enhanced, and showed higher c-Fos, intracellular Aβ expression, and axon sprouting. Granule cells only showed changes in synaptic properties, without intrinsic changes. The effects occurred before a memory task is affected.

**DISCUSSION:** Early electrophysiological and morphological alterations in Tg2576 MCs are consistent with enhanced excitability, suggesting an early role in DG hyperexcitability and AD pathophysiology.

**HIGHLIGHTS:**

∘ MCs from 1 month-old Tg2576 mice had increased spontaneous excitatory synaptic input.
∘ Tg2576 MCs had reduced spontaneous inhibitory synaptic input.
∘ Several intrinsic properties were abnormal in Tg2576 MCs.
∘ Tg2576 GCs had enhanced synaptic excitation but no changes in intrinsic properties.
∘ Tg2576 MCs exhibited high c-Fos expression, soluble Aβ and axonal sprouting.

## INTRODUCTION

Alzheimer’s disease (AD) is a neurodegenerative and progressive disorder characterized by memory impairment, and by the presence of amyloid β (Aβ) plaques and neurofibrillary tangles [1–4]. Despite extensive efforts, medications to treat AD are still needed. Several clinical studies suggest that treatment is most effective when started at early stages of AD. Therefore, it is relevant to understand the earliest contributing factors to AD.

Clinical findings suggest hyperexcitability is a characteristic of AD, often occurring at early stages, such as mild cognitive impairment (MCI) [5–11]. In patients, neuronal hyperexcitability is reflected by enhanced brain activity on fMRI when performing a memory-encoding task [12, 13], and by epileptiform activity and intermittent seizures [5-10, 14, 15]. Hyperexcitability is also present in mouse models of AD, which show intermittent seizures, interictal spikes, and high frequency oscillations, which are hallmarks of epilepsy [5, 16–25]. In addition, there is a reduction in the threshold for generating seizures in rodent models [26–28]. Interestingly, it was shown that reducing neuronal excitability can rescue the cognitive impairments in several mouse models [16, 29–34], and it also can reduce Aβ load, preventing the spread of Aβ pathology in the brain [21]. However, the underlying mechanisms of hyperexcitability and their importance to AD pathophysiology are not fully understood [7, 9, 14, 15].

An important area for studying hyperexcitability is the hippocampus because it is an early site for AD pathology [13] and is involved in seizure generation in temporal lobe epilepsy (TLE) [23, 25, 35, 36]. The hippocampal dentate gyrus (DG) has been suggested to be important in AD [37]. The DG is normally characterized by low excitability [38, 39], a product of the intrinsic properties of the main DG cell type, the granule cells (GCs) [38, 40, 41] and strong inhibition mediated by GABAergic interneurons [42–44]. As a result, DG can be considered as a gate that protects other hippocampal subfields from seizures. When impaired in AD, the normal role of the DG in spatial memory and other cognitive functions (such as pattern separation and novelty detection) are impaired [38, 45–48]. Moreover, GCs have been suggested to regulate seizures generation in TLE [38, 49–51].

The Tg2576 mice, an AD model that overexpresses a specific mutation in the amyloid precursor protein (APP; APPSwe), is characterized by a relatively slow progression of AD pathology [52–55], so one can study early stages. We previously showed that interictal spikes occur *in vivo* as early as 1 month of age, long before memory deficits and Aβ plaques [23]. When GCs were recorded in hippocampal slices, they showed increased excitatory synaptic input, as well as other changes, some of which might be compensatory [56]. Therefore, we asked if the increased synaptic input to GCs might be caused by hyperexcitability of one of their main excitatory inputs, the mossy cells (MCs).

MCs are glutamatergic neurons in the DG hilus that represent the primary excitatory input to the GCs proximal dendrites and constitute about 35% of all DG neurons [57–63]. MCs directly innervate GCs [64–66], but also DG interneurons [64], so MCs can regulate GCs by modulating the excitation/inhibition (E/I) balance. Although MCs have been studied in several diseases [47, 67–69], their contribution to AD has been rarely studied.

To study MCs and GCs at high resolution, we conducted whole cell recordings in hippocampal slices at 1 month of age and characterized spontaneous excitatory and inhibitory postsynaptic events (i.e., sEPSPs, sEPSCs and sIPSCs) and intrinsic properties (e.g., resting membrane potential, input resistance, etc.). Our results suggest that MCs have increased excitability at just 1 month of age. This was reflected in an increased E/I ratio of spontaneous synaptic events. In contrast, GCs did not show any changes in the E/I ratio, nor in their intrinsic properties. Furthermore, soluble Aβ is elevated in Tg2576 MCs, which also express more c-Fos protein than WT, confirming *in vitro* findings. Finally, a hippocampal-dependent behavioral task, novel object recognition, is not impaired at this age, suggesting changes in MCs occur before cognitive deficits.

## METHODS

### 1. Animals

All experimental procedures were carried out in accordance with the National Institutes of Health (NIH) guidelines and approved by the Institutional Animal Care and Use Committee (IACUC) at The Nathan Kline Institute. For Tg2576 mice expressing human APP_695_ with the Swedish (Lys670Arg, Met671Leu) mutations driven by the hamster prion protein promoter [52], male and female mice were obtained from crossing male heterozygous Tg2576 and female non-transgenic mice (C57BL6/SJL F1 hybrid, stock No. 100012, Jackson Labs). For Tg2576 and WT mice that expressed Cre-recombinase in cells with the dopaminergic D2 receptor (*Drd2*-cre), Tg2576^+/-^ males were crossed with *Drd2*-cre^+/-^ females (age range: 2-5 months). Breeders were fed a standard rodent chow (Purina 5008, W. F. Fisher). Offspring were housed with same-sex siblings, provided 2″× 2″ nestlets (W.F. Fisher), and fed rodent food after weaning (Purina 5001, W.F. Fisher). Food and water were available *ad libitum*, with a 12 h light-dark cycle. Genotypes for the APP_695_ gene and Cre gene were determined using standard protocols (New York University Mouse Genotyping Core) in Tg2576 mice and Tg2576 x *Drd2*-Cre mice.

### 2. Slice electrophysiology

#### 2.1 Slice preparation

Mice were deeply anesthetized by isoflurane (Pivetal, Aspen Veterinary Resources) inhalation, followed by an intraperitoneal (i.p.) injection of urethane (2.5 g/kg; Sigma-Aldrich) dissolved in 0.9% NaCl. Subsequently, mice were intracardially perfused with a cold (4°C) sucrose-based artificial cerebrospinal fluid (sucrose ACSF) containing (in mM): 90 sucrose, 2.5 KCl, 1.25 NaH_2_PO_4_, 4.5 MgSO_4_, 25.0 NaHCO_3_, 10.0 D-glucose, 80.0 NaCl, and 0.5 CaCl_2_, pH 7.4; constantly aerated with carbogen (95% O_2_, 5% CO_2_, All-Weld Products). The brain was quickly removed and dissected in sucrose ACSF at 4°C. Horizontal hippocampal slices (350 µm thick) were obtained using the same cold sucrose ACSF with a vibratome (VT1200S, Leica). Slices were immediately placed in a custom-made holding chamber containing sucrose ACSF at 30°C for 20 min and aerated with carbogen (95% O_2_, 5% CO_2_). Then, the slices were transferred and stored in the holding chamber containing normal recording ACSF (NaCl ACSF; containing NaCl instead of sucrose, in mM: 130 NaCl, 2.5 KCl, 1.25 NaH_2_PO_4_, 1 MgSO_4_, 25.0 NaHCO_3_, 10.0 D-glucose, and 2.4 CaCl_2_; pH 7.4) at room temperature for at least 60 min before recording.

#### 2.2 Whole-cell patch clamp recordings

##### 2.2.1 Recording acquisition

Slices were transferred to a recording chamber (RC-27LD, Warner) and perfused with NaCl ACSF at 6 mL/min with a peristaltic pump (Masterflex C/L, Cole-Parmer) and maintained at 32°C with a temperature controller (TC-324B, Warner) and in-line heater (SH-27B, Warner). For whole cell patch-clamp recordings, borosilicate glass capillaries (1.5 mm outer diameter and 0.86 mm inner diameter, Sutter Instruments) were pulled horizontally (P-97, Sutter) with a resistance of 4-9 MΩ. Current-clamp recordings were performed with a potassium (K^+^)-gluconate based intracellular solution containing (in mM; all reagents from Sigma-Aldrich): 130.0 K^+^-gluconate, 4.0 KCl, 2.0 NaCl, 10.0 HEPES, 0.2 EGTA, 4.0 Mg-ATP, 0.3 Na_2_-GTP, 14.0 Tris-phosphocreatine, and 0.5% biocytin (pH 7.3 and 295±5 mOsm). Voltage-clamp recordings were done using cesium (Cs^+^)-methanesulfonate intracellular solution of the following composition (in mM): 125 Cs^+^-methanesulfonate, 4 NaCl, 10 HEPES, 1 EGTA, 4 Mg-ATP, 0.3 Tris-GTP, 10 diTris-phosphocreatine, 5 QX-314 chloride, and 0.5% biocytin (pH 7.3 and 300±5 mOsm). Seal resistances were > 1 GΩ before breaking into whole cell configuration. All data were digitized (Digidata 1440A, Molecular Devices), amplified by a MultiClamp 700B amplifier (Molecular Devices), and low pass-filtered using a single-pole RC filter at 10 kHz. For the evaluation of synaptic activity, spontaneous events were recorded continuously for 3-5 min. Cells were included if access resistance did not change more than 20%. Data acquisition used pClamp software (v11.2, Molecular Devices) and data analysis was performed using Clampfit software (v11.2, Molecular Devices), as described further below.

##### 2.2.2 Spontaneous synaptic potentials and synaptic currents

Current-clamp recordings of spontaneous excitatory postsynaptic potentials (sEPSPs) were obtained from GCs using the K^+^-gluconate intracellular solution. Spontaneous EPSPs were evaluated at resting membrane potential (RMP). Voltage-clamp recordings of spontaneous excitatory and inhibitory postsynaptic currents (sEPSCs and sIPSCs, respectively) were performed in a different set of neurons, using the Cs^+^-methanesulfonate intracellular solution. Different holding potentials (HP) were used for isolating the different types of currents. To record sIPSCs, cells were voltage-clamped at a HP of +10 mV, and to evaluate AMPA- and NMDA-sEPSCs, cells were voltage-clamped at a HP −70 mV and −30 mV, respectively.

Detection of spontaneous events (sEPSPs, sEPSCs and sIPSCs) was performed off-line using MiniAnalysis 6.0 (Synaptosoft, Inc.) and the criteria described previously [56]. Briefly, sEPSPs, sEPSCs and sIPSCs were identified as events having a fast rise time, and were included if the maximum (peak) voltage (depolarizing, for EPSPs) or current (negative or positive, for EPSCs and IPSCs, respectively) was larger than 3 standard deviations (SD) from the root mean square (RMS) of the baseline noise [56, 70, 71]. The mean frequency and amplitude were calculated for all synaptic events over a 3 min recording period and are presented as mean ± SEM.

##### 2.2.3 Intrinsic properties

For intrinsic properties, current-clamp recordings were performed, and the different measurements are shown in Figure 2A. RMP was defined as the intracellular potential adjusted by the value recorded after withdrawing the microelectrode from the cell. To assess input resistance (R_in_), the steady-state voltage responses resulting from sub-threshold depolarizing and hyperpolarizing pulses (−30 pA to +30 pA in 5-10 pA current steps; 1 sec duration) were plotted against the amplitude of current injection and the slope of the linear fit between −5 pA to −30 pA was used to define R_in_ (Clampfit v. 11.2; Molecular Devices). Time constant (tau; τ) was determined from hyperpolarizing pulses (−20 pA) and defined as the time to reach 63% of the steady-state response. To determine action potential (AP) characteristics, depolarizing current pulses of +40 to +150 pA (10 pA steps; 1 sec duration) were delivered to elicit an AP close to its threshold in approximately 50% of trials, and a series of APs. We then used the AP analysis tool in Clampfit (pClamp software v11.2, Molecular Devices) to determine the characteristics of the APs. Threshold was defined as the membrane potential at which the AP initiated, considering the intersection of the AP rising phase and the slope leading up to its initiation. A three-point tangent slope vector was used to find the position in the initial region where the slope was at or above 10 V/sec. The mean AP peak amplitude was determined as the voltage difference from RMP to the AP peak. The time to the AP peak was the time from the start of the current step to the AP peak. The time to peak from threshold was measured as the period from the AP initiation to the AP peak. Half-width was defined as the period from the AP initiation to the point when the AP reached half of its peak amplitude. AP rising and decay slopes were defined by the maximum dv/dt of each AP phase, and the dv/dt ratio was defined as the ratio of rising/decay slopes. For the F-I curve, the mean firing frequency of APs generated by the application of depolarizing current steps (up to +150 pA in 10 pA steps, 1 sec duration) was evaluated. To quantify spike frequency adaptation (Supplementary Figures 1 and 2) we evaluated the time from one AP peak to the next for all the spike pairs in trains of 4 or 6-7 APs.

**Figure 1.**
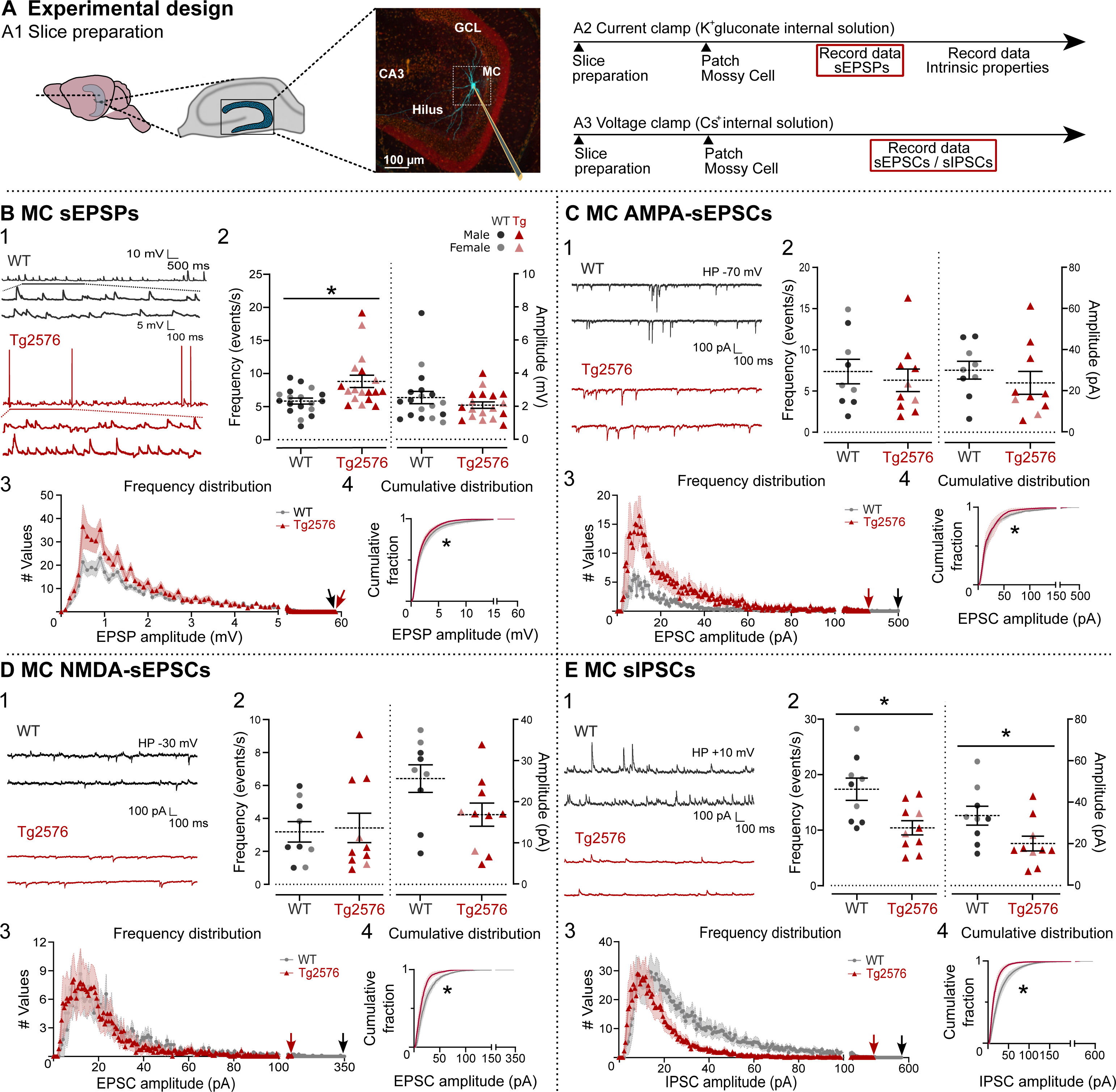
Tg2576 MCs exhibit increased excitatory and decreased inhibitory synaptic activity. **A.** 1. An illustration of the location of the hippocampus (left) and a cross-section through the hippocampus showing the DG (center). At right is a micrograph of the DG illustrating a microelectrode filled with biocytin that stained a MC (blue). 2-3. The timeline of the slice preparation and the recordings (2, current clamp; 3, voltage clamp) of spontaneous synaptic activity and intrinsic properties from MCs. **B.** 1. Representative traces showing typical sEPSPs, obtained from WT (black) and Tg2576 mice (red). 2. Quantification of sEPSP frequency and amplitude in WT and Tg2576 MCs. Mean sEPSP frequency was significantly greater in Tg2576 mice. 3. Frequency distributions of sEPSP amplitudes. Red and black arrows point to the maximums. 4. The cumulative representation of frequency distribution histograms revealed significant differences between WT and Tg2576 MCs. **C.** 1. Representative traces of AMPA-sEPSCs. 2. Quantification showed no significant differences in mean frequency or amplitude. 3. Frequency distributions suggested more small events in Tg2576 mice, and more large events in WT mice (compare arrows pointing to the maximums). 4. The cumulative distributions were significantly different in WT and Tg2576 mice, consistent with the frequency distributions. **D.** 1. Representative traces of NMDA-sEPSCs. 2. Quantification did not show differences in mean frequency or amplitude. 3. Frequency distributions showed more large events in WT mice. 4. Cumulative distributions were significantly different. **E.** 1. Representative traces of sIPSCs. 2. Quantification shows a significant reduction in mean sIPSC frequency and amplitude in Tg2576 mice. 3. Frequency distributions of events show significant differences. 4. The cumulative representation of frequency distribution histograms revealed significant differences between WT and Tg2576 MCs. For this and all other figures, data are represented as mean ± SEM. *p<0.05. Details of statistical comparisons are in the text and supplemental tables.

**Figure 2.**
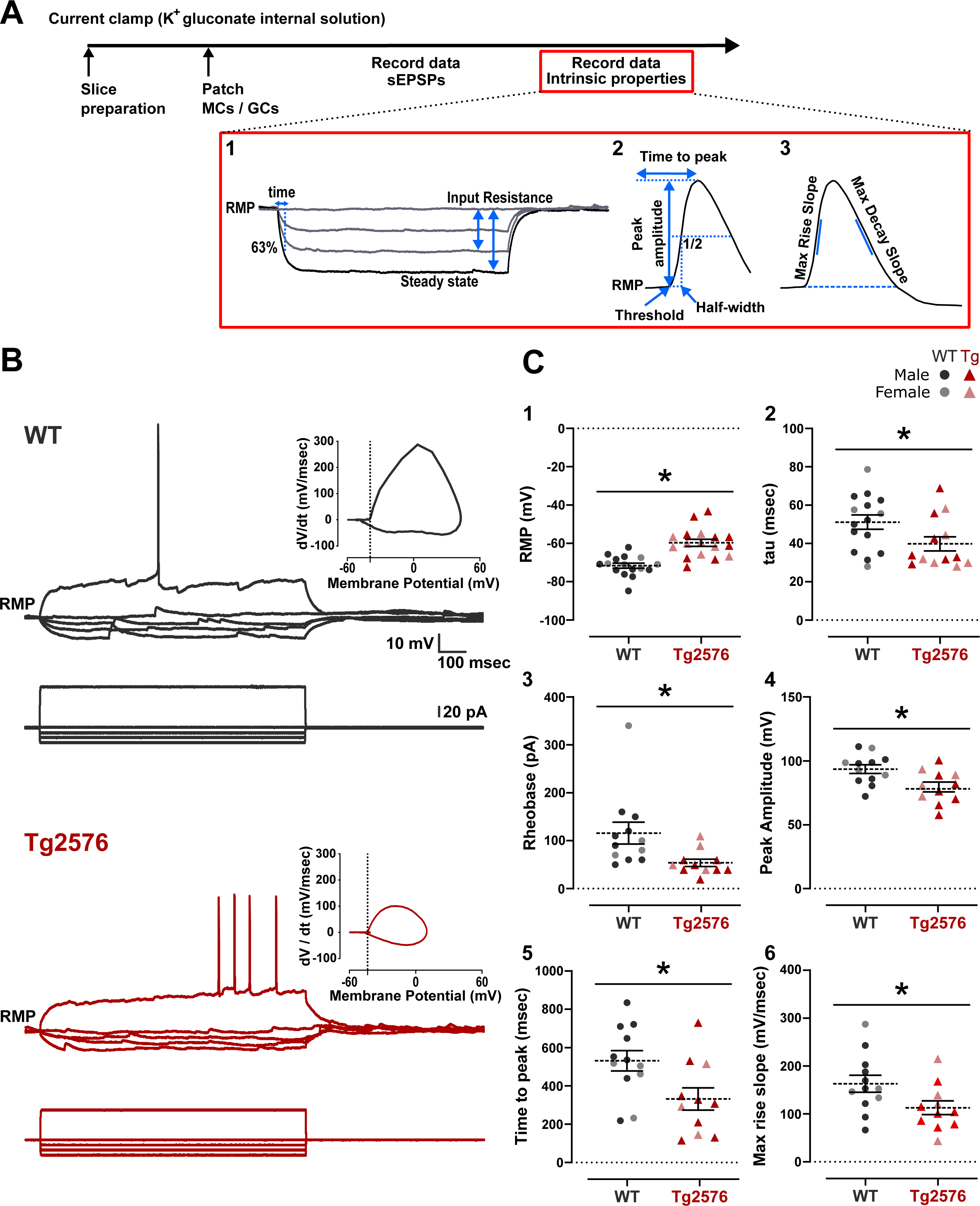
Tg2576 MC intrinsic properties of MCs suggest altered excitability. **A.** The timeline of the electrophysiological recordings and the analysis of intrinsic properties from MCs. 1. Measurements from responses to hyperpolarizing current steps. 2-3. Measurements of APs. **B.** Representative traces of membrane potential responses from WT (black) and Tg2576 MCs (red) to a positive (70 and 60 pA, respectively) and consecutive negative current steps from −10 to −30 pA using a 10 pA increment. The AP phase plot corresponding to the AP is shown in the inset above the traces. **C.** 1. Tg2576 MCs had significantly more depolarized RMPs than WT MCs. 2. Time constant was significantly shorter in Tg2576 mice. 3-4. Rheobase (3) and the time to peak of the AP (4) showed a significant reduction in Tg2576 mice. Together, data in 1-4 suggest enhanced excitability in Tg2576 mice. 5-6. AP peak amplitude (5) and maximum rising slope (6) was significantly reduced in Tg2576 mice, which would potentially decrease excitability.

**Figure 3.**
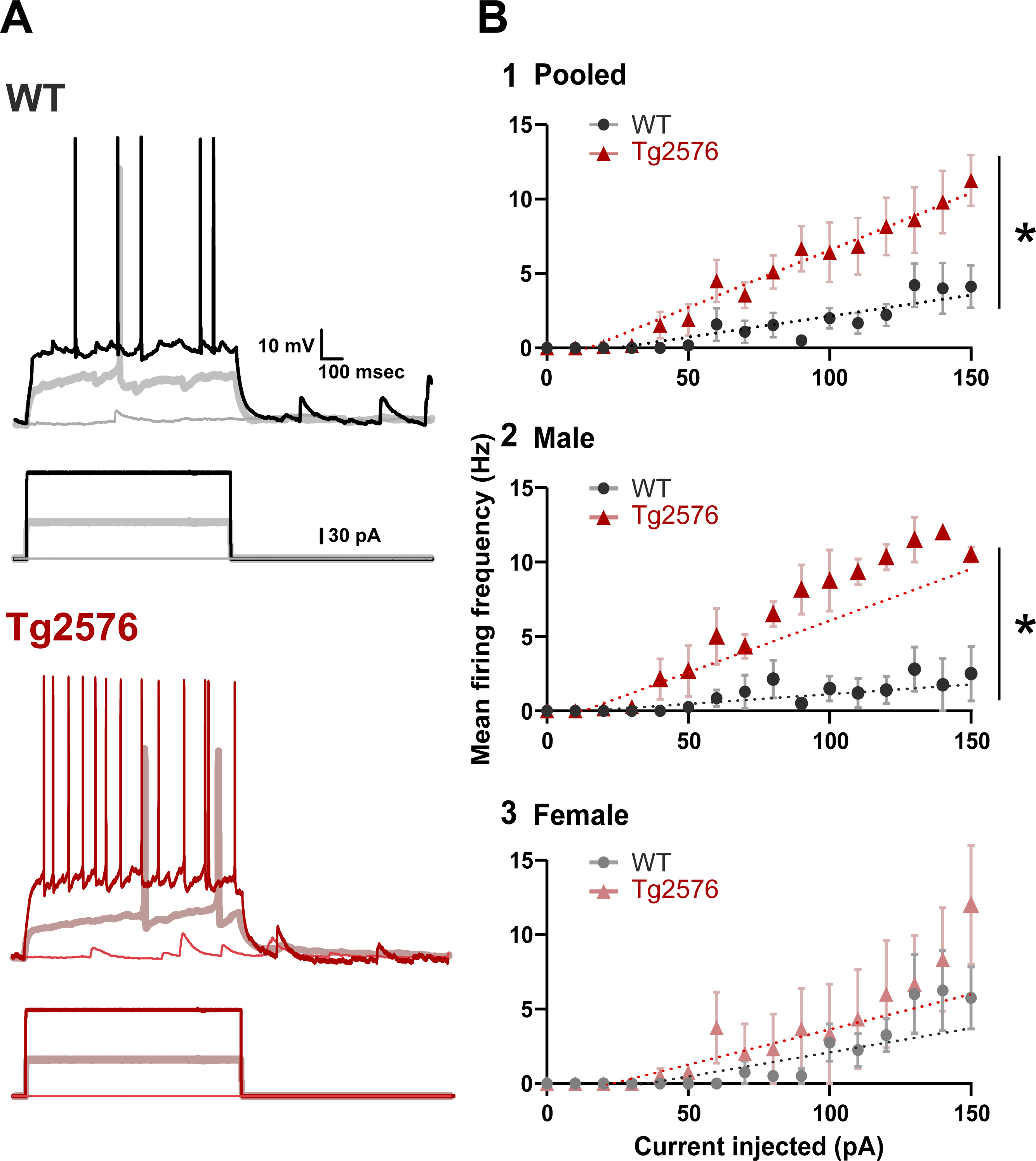
Enhanced F-I curve in Tg2576 MCs. **A.** Representative traces from WT (black traces) and Tg2576 MCs (red traces) of responses to positive current injection (0, 70 and 150 pA current steps). **B.** 1. The mean firing frequency was the average of all interspike intervals. The number of APs generated on each step was normalized by the time to obtain the mean firing frequency. The AP mean firing frequency was calculated for each response to a current step. Current steps were increased in 10 pA increments. Mean firing frequency of Tg2576 MCs was significantly increased compared to WT MCs. Both sexes were pooled. 2. Differences were significant in male mice. 3. Differences were not significant in females, which could be due to the variability of the female responses.

**Figure 4.**
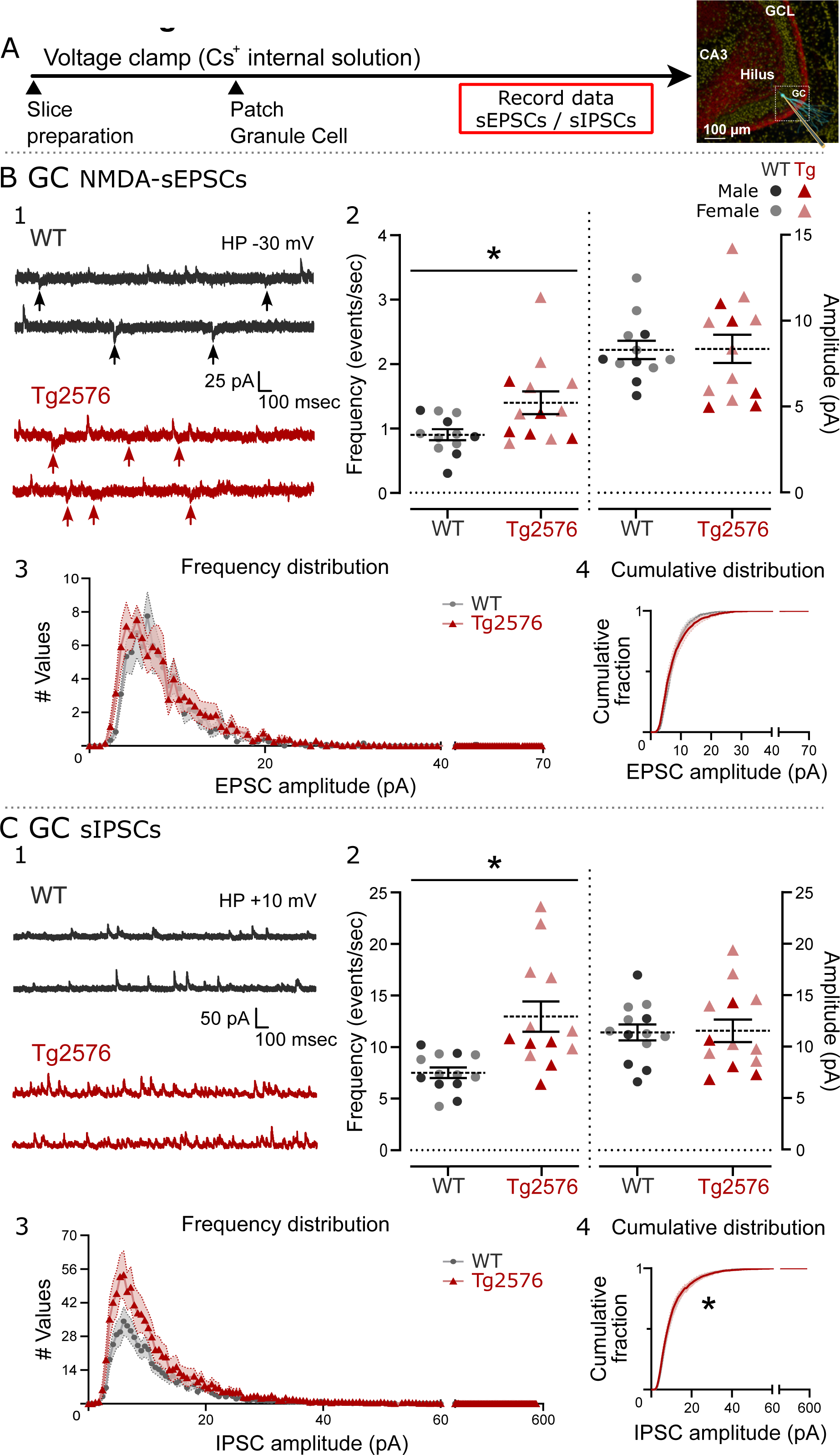
Tg2576 MC E/I balance suggests increased excitability. **A.** Increased E/I ratio in Tg2576 MCs compared to WT MCs. The ratio of sEPSCs (AMPA- and NMDA-sEPSCs) to sIPSCs is shown, based on mean frequency (1) or mean amplitude (2). **B.** 1-2. Charge transfer of excitatory currents (AMPA- and NMDA-sEPSCs) based on means (1) or all values (2). 3-4. Charge transfer of inhibitory currents (sIPSCs). 5. The E/I ratio for charge transfer based on mean values showed Tg2576 MCs had significantly increased charge transfer ratio.

### 3. Mossy cell and granule cell identification after electrophysiological recordings

After the completion of recordings, slices were immediately placed in a solution of 4% paraformaldehyde (PFA; Sigma-Aldrich) in phosphate buffer (PB; 0.1 M, pH 7.4) and kept in fixative at 4°C for 24-48 h. For biocytin processing, slices were washed in 0.1 M Tris buffer (TB; Sigma-Aldrich) and permeabilized with 1% Triton X-100 (TX, Sigma-Aldrich) in TB with continuous shaking on a rotator at room temperature for 20 min. Then, slices were incubated in a solution with 0.25% TX in TB for 1 h. Afterwards, slices were incubated in streptavidin-cyanine Cy2 conjugate (1:2000, Jackson ImmunoResearch) in 0.25% TX diluted in TB for 48 h at 4°C and protected from light. After this period, slices were washed in TB and exposed to a series of glycerol dilutions (25, 40, 55, 70, 85 and 90%; Sigma-Aldrich), immediately mounted on a 0.1% gelatin-coated slide surrounded by agar (4%) support, coverslipped using a mounting media solution of 0.5% N-propyl gallate (Sigma-Aldrich) and 90% glycerol in _dd_H_2_O, and the edges were sealed (CoverGrip, Biotium). Photomicrographs were acquired with a Zeiss LSM 880 laser scanning confocal microscope and Zen 3.0 software (Zeiss), with Plan-Apochromat 10×/0.45 M27 and Plan-Apochromat 20×/0.8 M27 objectives (Zeiss). All images were acquired at 16-bit depth with a frame size of 2048 x 2048 pixels.

### 4. Immunostaining for cFos, Aβ and MC axon identification

#### 4.1 Perfusion and sectioning

Mice were euthanized when their age was ∼1 month-old (33-38 days-old). Mice were anesthetized with isoflurane (Pivetal, Aspen Veterinary Resources), followed by urethane (2.5 g/kg; i.p.), and they were transcardially perfused with ∼20 mL of cold saline solution, followed by ∼20 mL of cold 4% PFA in 0.1 M PB using a peristaltic pump (Minipuls2, Gilson). The brain was quickly extracted and stored overnight in 4% PFA at 4°C. Afterwards, the left hemisphere was cut in the coronal plane and the right hemisphere was cut in the horizontal plane (50 μm-thick sections; Vibratome 3000, Ted Pella). The coronal and horizontal planes were used to optimize visualization of the sublamina of the DG molecular layer (ML), and in particular the MC axon terminal plexus in the inner molecular layer (IML). Sections were stored in 24-well tissue culture plates containing cryoprotectant solution (30% sucrose, 30% ethylene glycol in 0.1 M PB) at −20°C until use. In all cases, immunofluorescence staining was performed on free floating sections, and quantification was averaged from four horizontal sections, 600 µm apart, per subject.

#### 4.2 c-Fos immunostaining

Sections from WT and Tg2576 mice were processed together. Free-floating sections were washed in 0.1 M TB (3 washes, 10 min each) and subsequently incubated in two different solutions containing 0.25% TX in TB (TrisA), and 1% BSA and 0.25% TX in TB (TrisB), for 15 min each. Then sections were incubated in a blocking solution containing 5% normal goat serum (NGS; S-1000, Vector Laboratories), 1% BSA and 0.25% TX in TB for 60 min at RT. The primary antibody was a rabbit monoclonal anti-c-Fos antibody (Ab190289, Abcam) and was diluted in blocking solution (1:2000). Sections were incubated for 48 h at 4°C with continuous shaking. On the second day, sections were rinsed in TrisA and subsequently in TrisB for 15 min each, then incubated in goat anti-rabbit Alexa Fluor 488 secondary antibody (1:500; A11004, Invitrogen) diluted in TrisB and 3% normal goat serum (NGS; Vector Laboratories), for 2 h at RT, followed by a rinse in 0.1 M TB for 10 min. Next, sections were mounted on 0.1% gelatin-coated slides and allowed to dry for 1-2 h, and coverslipped with an antifade mounting medium containing DAPI (Vectashield HardSet H-1500, Vector Laboratories).

#### 4.3 Aβ immunostaining

For Aβ immunostaining, we used an antibody raised against the N-terminal fragment residues 1-12 of human Aβ (McSA1; MM0015p, Medimabs), which can detect oligomeric and highly aggregated insoluble forms of Aβ, as we previously reported [56]. Sections were treated with an antigen retrieval procedure where initially slices were washed in 0.1 M PB (3 washes, 10 min each), followed by an incubation period of 3 h in 0.1 M PB, pH 7.4, at 60°C to unmask the epitope of interest. Then, sections were rinsed in 0.1 M PB (3 washes, 10 min each) and incubated for 20 min in 0.5% TX diluted in 0.1 M PB, followed by incubation for 2 h in blocking solution containing 5% NGS (Vector Laboratories) and 1.5% Mouse-on-Mouse (MOM) blocking reagent (Vector Laboratories) in 0.1 M PB to block non-specific binding. Subsequently, sections were incubated for 48 h at 4°C in mouse monoclonal primary antibody to McSA1 (1:1000) in 3% NGS and 0.5% TX diluted in 0.1 M PB. On the second day, slices were washed in 0.1 M PB (3 washes, 10 min each) and incubated with goat anti-mouse IgG Alexa Fluor 488 secondary antibody (1:350, A11001, Invitrogen) for 2 h at RT. Finally, sections were rinsed in 0.1 M PB (3 washes, 10 min each), mounted on 0.1% gelatin-coated slides, allowed to dry, and then coverslipped with antifade mounting medium containing DAPI (Vectashield HardSet H-1500, Vector Laboratories).

#### 4.4 Calretinin immunostaining

To better visualize MC axons, we performed immunostaining against calretinin, a MC marker [72, 73]. We proceeded exactly as explained for c-Fos immunostaining but used a primary antibody to calretinin instead of c-Fos (1:2000; mouse anti-calretinin, 6B3, Swant). For calretinin and c-Fos double labelling, the secondary antibodies were chosen against the species of the primary antibody and with different Alexa Fluor color tags (1:500; goat anti-rabbit Alexa Fluor 488, A11004; goat anti-mouse Alexa Fluor 568; A11008, Invitrogen). The rest of the procedures were identical to 4.2.

#### 4.5 Image acquisition

Photomicrographs were acquired with a Zeiss LSM 880 laser scanning confocal microscope and Zen 3.0 software (Zeiss), with Plan-Apochromat 10×/0.45 M27 and Plan-Apochromat 20×/0.8 M27 objectives. All images were acquired at 16-bit depth with a frame size of 2048 x 2048 pixels. For high-resolution insets, the Plan-Apochromat 20×/0.8 M27 objective was used with a 2.4x digital zoom. Immunofluorescence was visualized with preconfigured excitation and emission wavelengths in the acquisition software for DAPI (Ex/EM 408/453 nm), Alexa 488/GFP (Ex/EM 488/535 nm), and Alexa 568/mCherry (Ex/Em 561/643 nm). Zen 3.2 Blue Edition software (Zeiss) was used offline to perform all quantification, as explained below, and to convert raw Zeiss image files (CZI format) into TIF format.

### 5. Quantification

#### 5.1 c-Fos-immunofluorescence (cFos-IF)

c-Fos-IF was quantified by defining two different regions of interest (ROI) encircling the hilus and the GC layer (GCL). The GCL was defined as the area where GC somata are packed together between hilus and the ML, and the hilus was defined as zone 4 of Amaral [61, 69] (Figure 6A). C-Fos-IF was quantified using Zen 3.2 Blue Edition software (Zeiss) on images acquired at 20x. C-Fos-IF was calculated by subtracting the background from the mean intensity. The background was defined as the average of two different ROIs with no cells (stratum lacunosum-moleculare and the outer third of the ML; OML). C-Fos-IF was normalized and is presented as the percentage of change from background (% of background).

**Figure 5.**
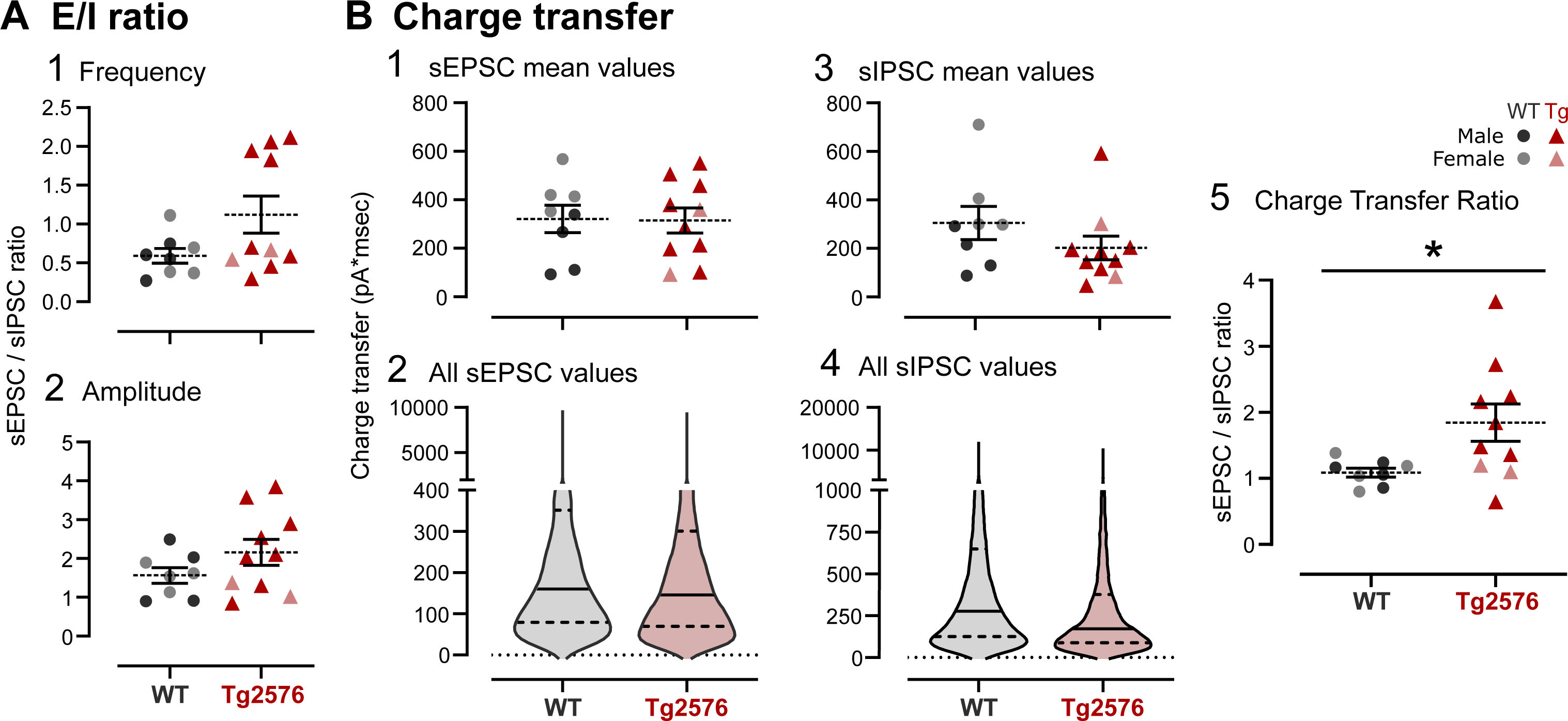
Increased spontaneous excitatory and inhibitory events in Tg2576 GCs. **A.** The timeline of the slice preparation and the electrophysiological recordings for spontaneous synaptic activity, which was similar for determination of intrinsic properties of GCs. To the right is a biocytin-filled GC (blue). **B.** GC NMDA-EPSCs. 1. Representative traces showing typical NMDA-sEPSCs from WT (black) and Tg2576 mice (red). 2. Tg2576 GCs had a higher frequency of sEPSCs but not amplitudes. 3. The frequency distributions and cumulative distributions did not show significant differences. **C.** GC sIPSCs. Representative traces of GC sIPSCs showed increased frequency but not amplitude of Tg2576 GCs. 3. Frequency distributions of sIPSCs amplitudes showed more small events in Tg2576 mice. 4. Cumulative distributions were significantly different.

**Figure 6.**
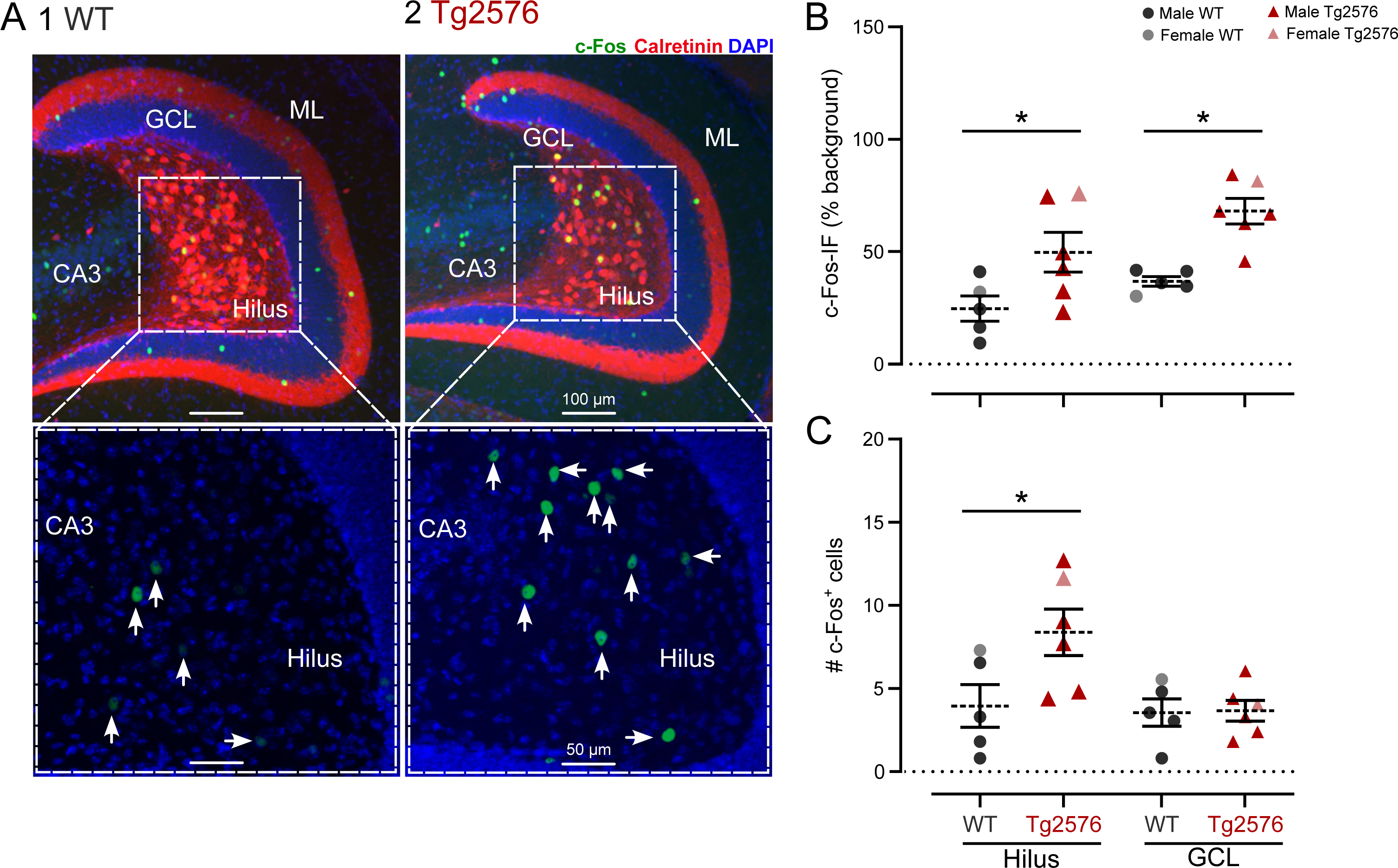
Increased c-Fos expression in the DG of Tg2576 mice suggests enhanced neuronal activity. **A.** Micrographs of horizontal sections from WT (1) and Tg2576 (2) DG showing immunofluorescence (IF) for c-Fos protein (green), calretinin (red) and DAPI (blue) in the hilus, granule cell layer (GCL), molecular layer (ML) and CA3. The boxed areas are expanded below and show hilar cells expressing c-Fos (green) that were counted based on thresholding. Arrows point to hilar cells considered immunolabeled. **B.** Quantification of c-Fos IF showed increased expression in Tg2576 hilus and GCL. **C.** There were significantly more c-Fos+ hilar cells in Tg2576 mice than WT mice. No significant differences were found for the number of GCs.

#### 5.2 c-Fos+ cell quantification

To quantify the number of c-Fos+ cells within each ROI, we used a thresholding method to count only the brightest cells and minimize background. From three to four sections were used per mouse, distributed within the horizontal sections that were cut. Hilar and GCL c-Fos+ cell bodies were quantified manually using the “events detection tool” in Zen 3.2 Blue Edition software (Zeiss).

#### 5.3 Aβ-immunofluorescence (Aβ-IF)

Aβ-IF quantification was performed as described for c-Fos-IF, where the two different ROI including the hilus and the GCL were defined, and the level of Aβ-IF was quantified using Zen 3.2 Blue Edition software (Zeiss) on 20x magnification images. Two different background values were evaluated from two locations without detectable Aβ-IF (in stratum lacunosum-moleculare and the OML) and averaged, which were subtracted from the values of intensity of the analyzed area for each section. Aβ-IF was normalized and presented in the results as the percentage of change from background (% of background).

#### 5.4 Quantification of MC axons

To quantify potential differences between the axons of Tg2576 and WT MCs, calretinin-mCherry+ immunofluorescence was evaluated in the IML (Figure 8). The ML was defined as the region between the fissure and GCL [46, 69] (Figure 8B, C). The IML, MML, and OML subdivisions of the ML were determined by dividing into equal thirds the total width of the ML, where the IML is closest to the GCL border and the OML ends at the hippocampal fissure. MC axon distribution was quantified in the upper blade (UB, suprapyramidal), crest (also called apex), and lower blade (LB, infrapyramidal) of the DG (Figure 8B), as previously defined [64].

**Figure 7.**
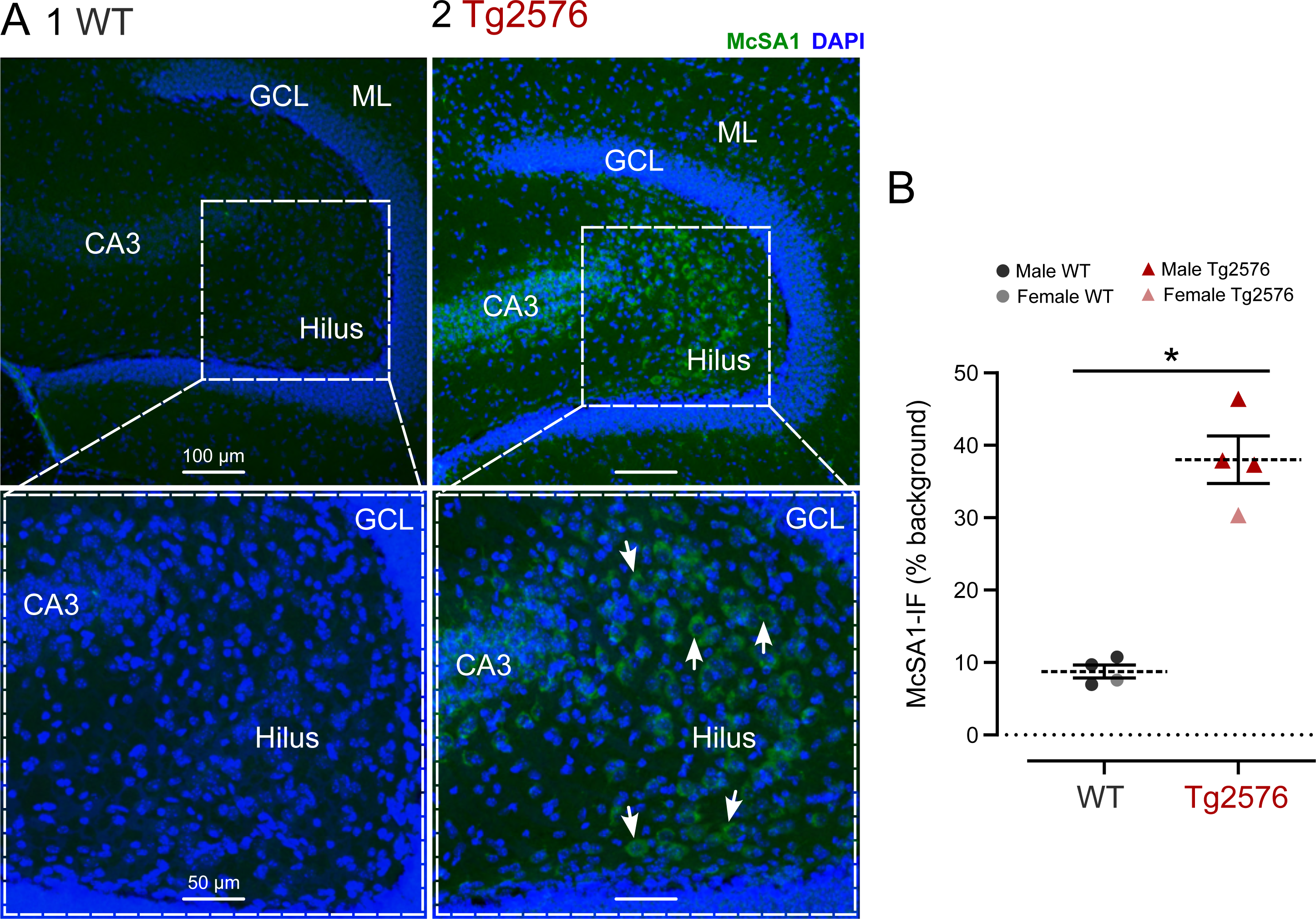
Increased Aβ expression in the DG of Tg2576 mice. **A.** Representative micrographs of immunohistochemical labeling of Aβ with the McSA1 antibody (green) and DAPI (blue) in horizontal sections for WT (1) and Tg2576 mice (2). Images show the hilus, granule cell layer (GCL), molecular layer (ML) and CA3. The boxed areas are shown at higher power below. Hilar cells expressing Aβ are shown in green and pointed with arrows. **B.** There was significantly more intracellular Aβ labeling with the McSA1 antibody for hilar cells from Tg2576 mice compared to WT.

**Figure 8.**
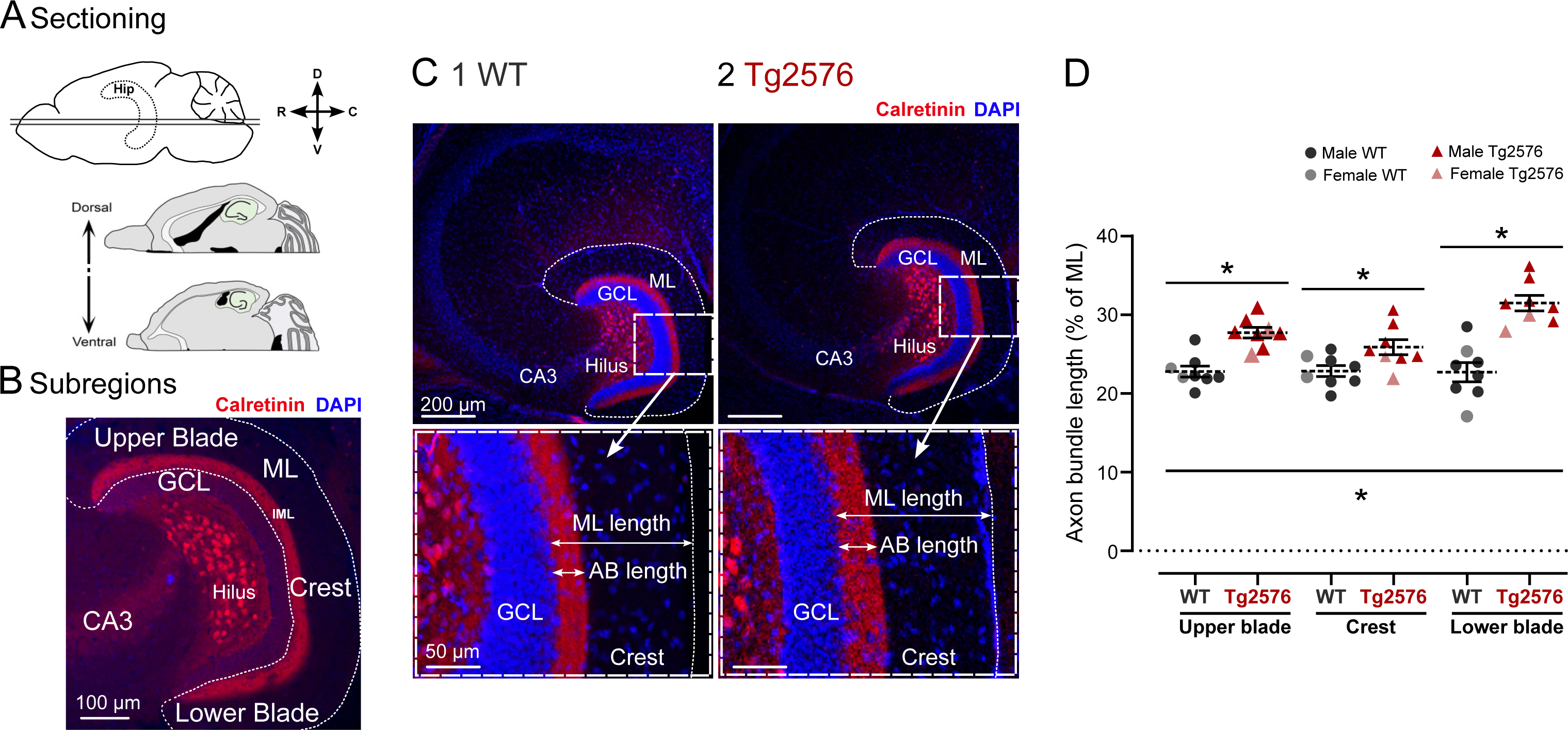
MC axons increase in Tg2576 mice. **A.** A diagram shows the plane that was used to generate horizontal sections selected from different parts of the septotemporal axis. **B.** The subregions of the DG are illustrated. Immunofluorescence for calretinin (red) and DAPI (blue) are shown. **C.** Micrographs of horizontal sections from WT (1) and Tg2576 mice (2). The boxed areas are expanded below and illustrate the measurements of the calretinin-stained MC axon plexus (axon bundle, AB length) and the ML length (horizontal arrows). **D.** Tg2576 mice showed an increase in the calretinin-stained plexus (quantified as the % of the ML).

The distance measurements were performed using the “distance tool” in the Zen 3.2 Blue Edition software (Zeiss). The length feature of the distance tool allows drawing parallel lines between two points to determine the length between them. Thus, one line from the GCL border to the edge of the mCherry+ axon terminal plexus (the dense band of mCherry+ puncta reflecting MC axon boutons) was drawn, along with one line from the GCL border to the end of the ML (Figure 8C1a and 8C2b). Distance measurements were normalized as the percentage of the total ML width (% of ML) by dividing our measurement of interest by the total width of the ML. All analyses were done using horizontal sections due to the ability to clearly show all the DG sub lamina, and a minimum of three relatively dorsal and three ventral sections were analyzed per mouse and averaged.

### 6. Novel object recognition (NOR) test

The NOR test has been shown to be a hippocampal-dependent task which involves the DG [74, 75]. For this test, mice were acclimated for three consecutive days (5 min each, Supplementary Figure 4A1), allowing every animal to freely explore a standard rat cage (26 cm wide x 40 cm long x 20 cm high) inside a larger cardboard box with no cover (40 cm wide x 60 cm long x 50 cm high). There were 3 pictures pasted to the walls of the box with different shapes (10-20 cm wide, 12-27 cm tall) and a combination of different colors (white, black, green, yellow, red) to provide a consistent context. Each picture was approximately centered on each wall and one edge touched the base of the box. All acclimation, training and testing sessions were performed between 10:00 a.m. − 12:00 p.m., and all equipment was cleaned using 70% ethanol between each session. On day 4, a training and testing session (5 min each) were conducted, with an interval between sessions of 1 h. Video recordings of all training and testing sessions were captured using a mobile phone camera with a 720p resolution and exported as H.264 MPEG-4/AVC videos. During the training session, mice were placed in the testing arena containing two identical objects (2 pineapple-like metal objects, 3 cm diameter x 6 cm height) centered along the shortest cage wall (Supplementary Figure 4A2) and allowed to explore them for 5 min. Afterwards, the mouse was removed and placed in its home cage for 1 h. For the testing session, one of the objects was replaced with a different one (a glass vial, 3 cm diameter x 6 cm height, filled with pieces of metal and salt to make it heavy; Supplementary Figure 4A3), and the mice were allowed to explore them for 5 min. Videos of the training and testing sessions were analyzed offline using ANY-maze video tracking system software (v7.3, Stoelting). Exploration was quantified as the time each mice spent exploring the objects, defining exploration as the nose pointed at the object and in a proximity of at least 2 cm of the object, and when animals spent time on top of the object, were looking down and sniffing it. The number of approaches to the object were also quantified. Exploration of the novel object during NOR testing was expected to be higher than 50% (as occurs for the training period) of the total time of object exploration, as this reflects animals can remember the objects seen in training (Supplementary Figure 4B).

### 7. Statistics

All statistical analyses were performed using Prism software (v. 10, GraphPad). For data that were normally distributed determined by the Shapiro-Wilk tests, parametric statistics were used. Otherwise, non-parametric tests were applied. For pairwise comparisons, homogeneity of variance was evaluated using the F test, and to compare multiple groups the Brown-Forsythe test or Bartlett’s test was used.

For parametric data, statistical significance between two groups was determined using the unpaired Student’s t-test, which was also used for data at two time points during the same event (e.g., train of APs). For non-parametric data, a Mann-Whitney *U* test was used for paired comparisons.

For analyzing cumulative distributions, we used the Kolmogorov-Smirnov (KS) test. For the frequency and current injected relationship (F-I curve), a linear correlation of the mean firing frequency vs. the current injected was determined, and then the Fisher z-transformation of the Pearson correlation coefficients was used to evaluate statistical differences between groups. A two-way repeated measures (RM) ANOVA (or mixed-effects model) followed by Šídák’s post-hoc test for the multiple comparisons was used to address the sequential APs of a train with genotype and current injected as factors. The same approach was used for analysis of spike frequency adaptation.

For evaluation of sex and genotype as factors, two-way ANOVAs were used. Šídák’s post-hoc test for the multiple comparisons was used. For the analysis of MC axonal distribution, comparisons between parametric data were performed using unpaired t-tests for two groups and ANOVAs for more than two groups. For non-parametric data, the Mann-Whitney and Kruskal-Wallis tests were used. When we asked whether the septotemporal area had an effect, two-way ANOVAs were used with location and genotype as factors. Šídák’s post-hoc test for the multiple comparisons was used.

All results are presented as the mean ± standard error of the mean (SEM), and statistical significance was achieved if the p value was <0.05 (denoted on all graphs by an asterisk).

## RESULTS

### 1. MC synaptic and intrinsic properties comparison between WT and Tg2576 slices

#### 1.1 Spontaneous synaptic events

There were 28 MCs from 22 Tg2576 mice and 28 MCs from 18 WT mice. For GCs, there were 25 cells from 16 Tg2576 mice and 24 GCs from 16 WT mice. No more than 3 cells (MCs or GCs) and 2-3 slices were used per mouse.

##### 1.1.1 Increased sEPSP frequency suggests enhanced excitatory activity in Tg2576 MCs

MCs were recorded using the current clamp configuration to assess sEPSPs (Figure 1). MCs from Tg2576 mice showed a significantly higher sEPSP mean frequency than WT mice (Tg2576: 8.82 ± 0.93 events/sec, WT: 5.84 ± 0.45; Mann-Whitney test, U=76, p=0.006; Figure 1B1, 1B2), as reflected by the representative traces in Figure 1B1. Despite this increase in frequency, Tg2576 MC sEPSPs did not show any differences in their mean amplitudes from WT (Mann-Whitney test, U=141, p=0.521; Figure 1B2). However, the frequency distribution of sEPSP amplitudes (Figure 1B3) showed a larger number of small events in Tg2576 MCs, as observed in the representative traces (Figure 1B1). Consequently, the cumulative distribution of these amplitudes was statistically different (Kolmogorov-Smirnov test, D=0.117, p=0.001; Figure 1B4).

##### 1.1.2 Spontaneous excitatory synaptic currents are increased in Tg2576 MCs

Next, AMPA receptor- and NMDA receptor-mediated sEPSCs were compared in WT and Tg2576 MCs in voltage clamp (Figure 1C, 1D). The experimental timeline for voltage clamp experiments (Figure 1A3) was analogous to current clamp, but different MCs were recorded due to the use of different intracellular solutions. As reported in our previous work [56], sEPSCs were identified as fast inward currents at −70 and −30 mV holding potentials, for AMPA- and NMDA-mediated currents, respectively, as shown in the representative traces in Figures 1C1 and 1D1. In our data set, we did not find significant differences between Tg2576 and WT MCs in AMPA-sEPSC mean frequency (unpaired t-test, t=0.5244, df=17, p=0.609; Figure 1C2), NMDA-sEPSC mean frequency (Mann-Whitney test, U=43, p=0.905; Figure 1D2), or AMPA-sEPSC mean amplitude (unpaired t-test, t=0.860, df=17, p=0.402; Figure 1C2), but there was a trend for reduced NMDA-sEPSC mean amplitude (unpaired t-test, t=2.021, df=17, p=0.059; Figure 1D2). However, the Tg2576 MC AMPA-sEPSC frequency distribution of amplitudes illustrated a greater number of small events, which led to a significant difference from WT MCs in the cumulative distribution (Kolmogorov-Smirnov test, D=0.367, p<0.001; Figure 1C4). Similarly, the NMDA-sEPSC frequency distribution of amplitudes showed a greater number of small sEPSCs in Tg2576 mice than WT (Kolmogorov-Smirnov test, D=0.428, p<0.001; Figure 1D4), possibly due to the trend for reduced amplitude evident in the means, as well as more large events in WT MCs. The greater number of large events in WT MCs is noted by a black arrow in Figure 1D3 (compared to red arrow marking the Tg2576 MC data). The significant genotype differences shown by cumulative distributions (but not means) may be because cumulative representations are more sensitive to the number of events of all amplitudes, whereas mean calculations pool all data.

##### 1.1.3 Inhibitory current events are mainly affected in Tg2576 MCs

We also evaluated whether the inhibitory currents of Tg2576 MCs (sIPSCs) differed from WT. For these experiments, the recorded MCs were the same as those used to record sEPSCs, but they were held at +10 mV holding potential, where sIPSCs were recorded as fast outward currents (Figure 1E1). In Tg2576 MCs, sIPSC mean frequency was significantly lower than WT (Tg2576: 10.42 ± 1.29 events/sec, WT: 17.38 ± 2.01; unpaired t-test, t=2.974, df=17, p=0.009; Figure 1E2). Furthermore, there was a significant reduction in sIPSC mean amplitude (Tg2576: 20.15 ± 3.52 pA, WT: 33.57 ± 4.56; unpaired t-test, t=2.356, df=17, p=0.031; Figure 1E2). Therefore, the frequency distribution of sIPSC amplitudes in Tg2576 MCs showed a reduced number of events (∼20-80 pA; Figure 1E3), and a significant difference in the cumulative distribution (Kolmogorov-Smirnov test, D=0.088, p=0.012; Figure 1E4).

Taken together, data from current and voltage clamp suggest excitation was increased in Tg2576 MCs, probably because of an increase in small EPSP/AMPA-EPSCs in Tg2576 mice, and a reduced number of large EPSCs compared with WT. Furthermore, there was a significant decrease in inhibitory synaptic input to MCs of Tg2576 mice. Therefore, the data suggest an increased E/I balance in Tg2576 MCs, which could contribute to a very early increase in MC excitability in Tg2576 mice.

#### 1.2. MC intrinsic properties in Tg2576 mice reflect enhanced excitability

Next, we examined MC intrinsic properties to determine whether MCs are intrinsically more excitable in Tg2576 mice. We evaluated RMP, R_in_, tau, and characteristics of APs. The AP characteristics were amplitude, measurements of duration (time to peak, half-width), threshold, maximum rate of rise, maximum rate of decay, dv/dt ratio (maximum rate of rise / maximum rate of decay), and rheobase (see Methods and Figure 2A).

The RMP was more depolarized in Tg2576 MCs relative to WT mice (−59.70 ± 1.85 mV vs −71.66 ± 1.33, respectively; unpaired t-test, t=5.197, df=31, p<0.001; Figure 2C1). Tau was shorter in Tg2576 than WT mice (39.81 ± 3.69 msec vs. 51.14 ± 3.74, respectively; unpaired t-test, t=2.142, df=26, p=0.042; Figure 2C2). Representative APs at threshold and their phase plots are shown in Figure 2B. Tg2576 MCs showed a significant reduction in the amount of current necessary to elicit an AP, or rheobase, in comparison with WT MCs (54.55 ± 7.67 pA vs. 115.8 ± 22.75, respectively; Mann-Whitney test, U=18, p=0.002; Figure 2C3). AP amplitude was reduced in Tg2576 mice relative to WT (79.57 ± 3.86 mV vs. 93.50 ± 3.67, respectively; unpaired t-test, t=2.731, df=21, p=0.013; Figure 2C4), as shown in the representative traces (Figure 2B). The time to the AP peak was shorter in Tg2576 than WT mice (329.20 ± 58.30 msec vs. 531.50 ± 53.30; unpaired t-test, t=2.565, df=21, p=0.018; Figure 2C5), suggesting Tg2576 MCs fire easier than WT. Finally, Tg2576 MC APs have a slower maximum rising slope than WT (113.00 ± 14.36 mV/msec vs. 163.20 ± 17.70; unpaired t-test, t=2.175, df=21, p=0.041; Figure 2C6). Tg2576 MCs did not differ from WT mice in the R_in_, threshold, time to peak from threshold, half-width, maximum rate of decay and the dv/dt ratio (Supplementary Table 1).

#### 1.3. Tg2576 MCs exhibit enhanced firing behavior

To assess firing behavior, we first evaluated MC AP firing induced by current injection (representative traces in Figure 3A). The relationship between mean firing frequency and injected current (F-I curve) is shown in Figure 3B. In comparison with WT, Tg2576 MCs fired more APs for any injected current step, which is depicted in the separation of the two genotypes by linear regression (Figure 3B1). These differences were significant when comparing the correlation factor for both genotypes (Fisher z-value for Tg2576 and WT, rz=2.417 and 1.487, respectively; z-score=-2.460). A two-way RMANOVA (mixed effects model) showed significant differences in genotype (F(1,21)=12.88; p=0.002). The other main factor, intensity of current, was also significantly different since current was increased sequentially. Thus, Tg2576 MCs were more inherently excitable compared with WT.

We also evaluated spike frequency adaptation in both genotypes (Supplementary Figure 1). To quantify adaptation, the interspike intervals (measured as the time from one AP peak to the next, or peak to peak time) were compared with trains of 4 APs (3 spike-pairs, SP; Supplementary Figure 1A) or longer AP trains (6 SPs; Supplementary Figure 1B). In general, MCs from Tg2576 and WT mice seemed to have weak adaptation, because there were similar interspike intervals during the train. A two-way RMANOVA (mixed-effects model) showed no effect of genotype for the 3 AP train (p=0.119) and 6 AP train (p=0.769; Supplementary Figure 1A1, 1B1). We did not find any significant effect of genotype when the first and third SP were compared in each of the trains with 3 SPs for Tg2576 and WT MCs (p=0.791 and p=0.999, Šídák’s multiple comparisons test, respectively), or when we compared the first and sixth SP using longer trains (p=0.933 and p>0.999, Šídák’s multiple comparisons test, respectively).

The data suggest that MCs have weak spike frequency adaptation in both genotypes. This lack of adaptation has been suggested to occur because MCs have a great deal of spontaneous input which leads to high variability in their AP firing pattern [58].

#### 1.4. Tg2576 MCs have an increased E/I ratio

The E/I ratio of sEPSCs and sIPSCs of Tg2576 MCs was not significantly different from WT for their frequency and amplitude (Mann-Whitney test, U=23, p=0.146, and unpaired t-test, t=1.431, df=16, p=0.172, respectively; Figure 4A1, 4A2). When we analyzed the charge transfer of excitatory and inhibitory events, there were no significant changes for sEPSCs (unpaired t-test, t=0.070, df=16, p=0.945; Figure 4B1), or sIPSCs (Mann-Whitney test, U=24, p=0.173; Figure 4B3). However, estimation of the E/I ratio based on these values showed a significant increase in Tg2576 mice compared to WT mice (1.84 ± 0.28 vs. 1.09 ± 0.07, respectively; Mann-Whitney test; U=5, p=0.027; Figure 4B5). The reason could be a small increase in EPSP/EPSCs and decrease in IPSCs, although the means were not different.

Taken together, the data suggest a very early increased excitability of Tg2576 MCs. The increased excitability is likely to be due to increased sEPSPs/Cs, decreased sIPSCs, and altered intrinsic properties.

#### 1.5. Comparison of sex in WT and Tg2576 MCs

Female and male mice were compared for all measurements described above. However, females were studied without knowing the stage of their estrous cycle. Therefore, the combination of physiological changes dependent on different cycle stages were likely to be underrepresented.

Regarding MC synaptic events, when data were analyzed using a two-way ANOVA with sex and genotype as main factors, we did not find any significant sex differences, but genotype was a significant factor (Supplementary Table 2).

When analyzing MC intrinsic properties with a two-way ANOVA with sex and genotype as main factors, we did not find any significant sex differences, but genotype was a significant factor for RMP, tau, rheobase, AP peak amplitude, time to peak and the dv/dt ratio (Supplementary Table 3).

### 2. GC synaptic and intrinsic properties in WT and Tg2576 mice

#### 2.1. Excitatory and inhibitory synaptic events are increased in frequency in Tg2576 GCs

Synaptic events and intrinsic properties were compared in WT and Tg2576 GCs following the same protocol described for MCs (Figure 5A). Tg2576 GC sEPSPs were similar to WT (Mann-Whitney test, p=0.487, for frequency, and unpaired t-test, p=0.996, for amplitude; Supplementary table 4), and so were AMPA-sEPSCs (unpaired t-test, p=0.534, for frequency, and unpaired t-test, p=0.471, for amplitude; Supplementary table 4). However, Tg2576 GCs showed a significantly higher frequency of NMDA-sEPSCs (1.40 ± 0.18 events/sec vs. 0.90 ± 0.09 for Tg2576 and WT, respectively; unpaired t-test, t=2.543, df=17.19, p=0.021; Figure 5B1, 5B2) and sIPSCs (12.97 ± 1.47 events/sec vs. 7.51 ± 0.51 for Tg2576 and WT, respectively; unpaired t-test, t=3.513, df=14.81, p=0.003; Figure 5C1, 5C2) than WT mice. These differences are exemplified in representative traces in Figure 5B1 and 5C1. The frequency distribution of sIPSC amplitudes showed a larger number of small events in Tg2576 GCs (Figure 5C3), and statistically different cumulative distributions (Kolmogorov-Smirnov test, D=0.352; p<0.001; Figure 5C4). These data suggest that the presynaptic inputs to GCs may be enhanced, and MCs may play a role as they can excite GCs directly and inhibit them indirectly by exciting interneurons [61, 64]. Therefore, increased MC excitability would increase EPSCs and IPSCs to GCs leading to few net effects. However, why NMDA-sEPSCs were increased but not AMPA-sEPSCs is not clear.

#### 2.2. No changes in GC intrinsic properties occur at very early ages in Tg2576 mice

GC intrinsic properties showed changes in 2-3 month-old Tg2576 mice [56], but at 1 month of age, GCs did not have any significant difference from WT in RMP, tau, rheobase, threshold, peak amplitude, time to peak, maximum rise slope, maximum decay slope and the dv/dt ratio (Supplementary Table 5). However, the AP half-width tended to increase (unpaired t-test, p=0.052; Supplementary Table 5), while Rin tended to reduce (unpaired t-test, p=0.079; Supplementary Table 5). These data suggest that GCs show synaptic changes before intrinsic properties change in Tg2576 mice. The latter might be compensatory because the changes that were found in the older mice mostly decreased GC excitability, as we previously reported [56].

#### 2.3. Tg2576 GC firing behavior is not different from WT at 1 month of age

To assess GC firing behavior, we evaluated the mean AP frequency induced by current steps (F-I curve; Supplementary Figure 2). In comparison with WT, Tg2576 GCs did not have any significant difference (two-way ANOVA, mixed effects analysis, p=0.172; Supplementary Figure 2A). Thus, the linear regressions for the two genotypes showed no significant differences (Fisher z-value for Tg2576 and WT, rz=2.247 and 2.359, respectively, z-score=0.297; Supplementary Figure 2A).

When we compared the interval between SPs for short and long trains, Tg2576 GCs did not differ from WT mice (two-way RMANOVAs, p=0.515 for 3 SPs, and p=0.060 for 6 SPs; Supplementary Figure 2B1, 2C1). When the first and sixth SPs were examined specifically in the longer AP trains, we found significant differences in Tg2576 GCs, suggesting adaptation occurred (Wilcoxon test, W=21, p=0.031; Supplementary Figure 2C2). Similarly, WT GCs also showed adaptation (Paired t-test, t=4.229, df=7, p=0.004; Supplementary Figure 2C2). These data suggest that GCs from both genotypes were able to adapt their firing pattern. There were no differences between genotypes (unpaired t-test, t=0.786, df=13, p=0.223, for SP1, and Mann-Whitney test, U= 14, p=0.114, for SP6; Supplementary Figure 2C2).

These datasets suggest the firing behavior of Tg2576 GCs is not affected at 1 month of age, which is consistent with the lack of significant changes in their intrinsic properties.

#### 2.4. Tg2576 GCs have a normal E/I ratio

Consistent with an increased NMDA-excitation and increased inhibition input frequency, we found the Tg2576 E/I ratio for frequency (Mann-Whitney test, U=61, p=0.243; Supplementary Figure 3A1) and amplitude (unpaired t-test, t=1.007, df=24, p=0.324; Supplementary Figure 3A2) was not significantly different from WT. Similarly, the charge transfer for excitatory and inhibitory events did not show differences between genotypes (Mann-Whitney test, U= 42, p=0.052, and Mann-Whitney test, U=65, p=0.336, respectively; Supplementary Figure 3B1, 3B2) and the E/I ratio of the charge transfer was not significantly different either (Mann-Whitney test, U= 53, p=0.186; Supplementary Figure 3B3). Thus, these data suggest GCs at 1 month of age have increased NMDA-sEPSCs and increased sIPSCs which produce no significant change in E/I ratio.

#### 2.5. Sex differences in WT and Tg2576 GCs

No significant differences were found between WT and Tg2576 male and female mice in the sEPSP frequency or amplitude (Supplementary Table 6). When we evaluated the differences between excitatory currents, the two-way ANOVA showed a main effect of sex (F(1,22)=4.506, p=0.045; Supplementary Table 6). *Post-hoc* tests showed that WT females had greater AMPA-EPSC amplitudes than WT males (Šídák’s multiple comparisons test, p=0.025). When sIPSC frequency data were analyzed using a two-way ANOVA, we found significant effect of sex (F(1,22)=5.097, p=0.034; Supplementary Table 6) and genotype (F(1,22)=11.89, p=0.002; Supplementary Table 6). *Post-hoc* test showed that Tg2576 females had greater IPSC frequencies than Tg2576 males (Šídák’s multiple comparisons test, p=0.030).

Interestingly, although we did not find any significant differences when we evaluated GC intrinsic properties with sexes pooled, when separating them by sex we did (Supplementary Table 7). When analyzing these data with a two-way ANOVA with sex and genotype as main factors, we found significant differences of sex in the Rin (F(1,16)=6.423, p=0.022), rheobase (F(1,16)=7.515, p=0.015), half-width (F(1,16)=13.67, p=0.002), and maximum decay slope (F(1,16)=8.605, p=0.010); and genotype only the half-width (F(1,16)=9.949, p=0.006) and maximum decay slope (F(1,16)=6.750, p=0.019). Regarding sex, *post-hoc* tests showed that Tg2576 females had a shorter half-width than Tg2576 males (Šídák’s multiple comparisons test, p<0.001), and WT females had a significantly increased maximum decay slope than WT males (Šídák’s multiple comparisons test, p=0.043). Regarding genotype, *post-hoc* tests only showed that Tg2576 males had a larger half-width than WT males (Šídák’s multiple comparisons test, p<0.001).

### 3. MC and GC morphology and location

To confirm cell identity, recorded MCs and GCs were filled with biocytin by diffusion from the patch electrode into the cell (see Methods). A randomly selected group of slices with recorded cells were processed to visualize biocytin (see Methods). An example of a filled MC is shown in Figure 1A1, and a recorded GC is shown in Figure 5A. For MCs, we identified them as large (∼25 µm) cells located in the hilus of DG, with a morphology consistent with a normal adult MC [61]. MC somata were multipolar and had dendrite branches extended throughout the hilar area as previously described [58]. The presence of thorny excrescences was specific for MCs, as previously reported [61, 69].

MCs were sampled from horizontal sections that were divided into relatively dorsal and ventral levels as shown in Figure 8A. For dorsal sections from WT mice, 13 MCs were sampled, and 3 MCs were close (within ∼100 µm) to the UB, 5 were close to the crest, 1 MC was close to LB, and 4 MCs were in the middle portion of the hilus (close to the border of the hilus with CA3c). From relatively ventral slices of WT mice, 11 MCs were sampled, and 4 MCs were close to the crest, 3 MCs close to the LB, and 4 MCs in the middle portion of the hilus. For Tg2576 mice, 8 MCs were sampled from relatively dorsal slices, and 1 MC was close to the UB, 1 MC close to the crest, 1 MC close to the LB, and 5 MCs in the middle portion of the hilus. From relatively ventral slices, 17 MCs were sampled, and 3 MCs were close to the UB, 7 MCs close to the crest, 1 MC close to the LB, and 6 MCs in the middle portion of the hilus. In this random sample of MCs, the relative numbers of MCs in dorsal vs ventral locations were not significantly different between genotypes (Fisher’s exact test, p=0.156). The relative numbers of MCs located near the UB, crest, LB, and middle of the hilus were not significantly different for the dorsal WT vs dorsal Tg2576 mice (Fisher’s exact test, p=0.504) or ventral WT vs. ventral Tg2576 mice (Fisher’s exact test, p=0.329). Thus, position did not appear to be a contributing factor to increased excitability of Tg2576 MCs.

For GCs, the somata was round or oval, ∼10 µm long, and located near the center of GCL, as for our previous study [56]. GCs dendrites extended throughout the ML and contained numerous spines, indicating they were mature GCs [46, 76]. Some GCs were sampled from different portions of the GCL to identify their relative position. For WT, 15 GCs were sampled from relatively dorsal slices, where 4 GCs were in the UB and 11 GCs in the crest. From relatively ventral slices, 8 GCs were sampled, where 1 GC was in the UB and 7 GCs in the crest. For Tg2576 mice, 4 GCs were sampled from relatively dorsal slices, where 1 GC was in the UB, 2 GCs in the crest and 1 GC in the LB. From relatively ventral slices, 18 GCs were sampled, where 2 GCs were in the UB, 7 GCs in the crest and 1 GC in the LB. In this random sample of GCs, there were no significant differences in distributions of GCs in different parts of the GCL (dorsal WT vs. dorsal Tg2576, Fisher’s exact test, p=0.245) or ventral WT vs. ventral Tg2576 (p>0.999), suggesting that the location in the GCL could not explain any genotypic differences.

### 4. Immunohistochemical analysis of c-Fos-IF and c-Fos+ MCs

#### 4.1 Increased c-Fos protein expression in Tg2576 hilar cells

To validate the results regarding the increased activity of Tg2576 MCs in slices, we evaluated the expression of a commonly used marker of neuronal activity, c-Fos (Figure 6), using similar methods to past studies [75, 77–79].

Our results showed that there was significantly greater c-Fos-IF in the DG of Tg2576 mice relative to WT (Figure 6). The hilar area of Tg2576 mice, where MCs are located, had a significant increase in c-Fos-IF (49.75 ± 8.86% vs. 24.65 ± 5.59; unpaired t-test, t=2.279, df=9; p=0.048; Figure 6A2, 6B). Similarly, the Tg2576 GCL had greater c-Fos-IF than WT (68.05 ± 5.68% vs. 36.79 ± 2.16; unpaired t-test, t=4.751, df=9; p=0.001; Figure 6A2, 6B). This increased expression suggests enhanced neuronal activity of the cells located in those areas.

#### 4.2. Increased numbers of c-Fos+ MCs correlate with the electrophysiological hyperactivity in Tg2576 mice

In order to know whether the number of cells that expressed c-Fos protein differed between genotypes, we counted the number of bright cells using a threshold. We set threshold at a percentage of the maximum IF so that the brightest cells were easily distinguished from background (see insets in Figures 6A1, 6A2). The results showed significantly more c-Fos+ cells in the hilus of Tg2576 mice relative to WT (8.34 ± 1.40 cells vs. 3.90 ± 1.28; unpaired t-test, t=2.290, df=9, p=0.048; Figure 6C), but not in the GCL (unpaired t-test, t=0.110, df=9, p=0.915; Figure 6C). Thus, there was not only an increased general fluorescence of hilar c-Fos in Tg2576 mice, but also an enhanced number of c-Fos-expressing hilar cells in Tg2576 mice. In a randomly selected subset of 59 c-Fos+ cells, the majority (81.4%) were double-labeled with calretinin, and therefore were MCs. These data are consistent with prior studies showing strong c-Fos expression of hilar cells from horizontal sections in mice and rats [75, 77]. They provide additional *in vivo* support for the idea based on *in vitro* electrophysiology that MCs have increased excitability in Tg2576 mice.

### 5. Robust Aβ expression in the hilus of Tg2576 mice

Because we found increased excitability of MCs in slices, we asked if Aβ was expressed in MCs. For this question we used the antibody McSA1, which labels plaques as well as intracellular (soluble) Aβ as we reported before [56]. Representative micrographs of the double labeling of McSA1 and DAPI are presented in Figure 7A. The McSA1 fluorescence in Tg2576 mice showed that intracellular Aβ was present in hilar cells. We observed there was a very low immunofluorescence in the WT mice. Therefore, Tg2576 mice had significantly greater immunofluorescence (38.00 ± 3.29% vs. 8.74 ± 0.89; unpaired t-test, t=8.595, df=6, p<0.001; Figure 7A2, 7B).

These data suggest that there is a robust and increased amount of soluble oligomeric Aβ at 1 month of age in Tg2576 hilus, which is consistent with our previous findings where soluble Aβ levels were elevated at 3 months of age [56, 80]. Therefore, it is possible that the increased excitability of Tg2576 MCs may be due to soluble intracellular Aβ, as has been reported by other studies showing Aβ can lead to altered cellular and synaptic function in other types of neurons [81–89].

### 6. Immunohistochemical identification of the MC axon distribution

#### 6.1 MC axon distribution in the ML of DG is expanded in Tg2576 mice

To quantify the extent of the axon plexus of MCs in the ML, we used calretinin to stain MC axons. First, we identified the two edges of the axon plexus: 1) the border with the GCL, and 2) the border with the ML (Figure 8B, 8C). We found that in Tg2576 mice, calretinin was a greater percentage of the ML compared with WT in the UB (27.72 ± 0.69% vs. 22.78 ± 0.70, respectively; unpaired t-test, t=5.122, df=14, p<0.001; Figure 8D), crest (25.89 ± 0.97% vs. 22.85 ± 0.70, respectively; unpaired t-test, t=2.548, df=14, p=0.023; Figure 8D), and LB (31.48 ± 0.98% vs. 22.70 ± 1.23, respectively; unpaired t-test, t=5.582, df=14, p<0.001; Figure 8D). Two-way ANOVA analysis revealed a significant main effect of genotype (F(1,42)=9678, p<0.001; Figure 8D), and also for the DG blade location (F(2,42)=5762, p<0.001; Figure 8D). Multiple comparisons test showed significant differences between Tg2576 and WT mice in the UB (p<0.001), crest (p<0.001) and LB (p<0.001).

### 7. NOR test

Finally, we asked if increased excitability of MCs influenced behavior of Tg2576 mice at 1 month of age. We chose the NOR test because it has been reported that the DG influences this task [74, 75]. The results of the NOR task are presented in the supplementary Figure 4. During the training session (Supplementary Figure 4A2, 4B), neither WT (paired t-test, t=0.195, df=3, p=0.858) nor Tg2576 mice (paired t-test, t=1.354, df=4, p=0.247) showed any preference (exploration time) for one object over the other, i.e., novel object exploration during training approached 50% in both groups (Supplementary Figure 4B), independent of genotype. This effect was also evident in the discrimination index (0.012 ± 0.058 and 0.013 ± 0.059 for Tg2576 and WT, respectively; Mann-Whitney test, U=9, p=0.849; Supplementary Figure 4C1) where values close to zero suggest no differences in the amount of time the mice explore both objects.

For the NOR testing period (Supplementary Figure 4A3), the time WT and Tg2576 mice explored the new object was greater than the familiar object (Supplementary Figure 4B). We observed that Tg2576 (61.33 ± 2.92% vs. 50.63 ± 2.89 for Test and Train periods, respectively; paired t-test, t=2.951, df=4, p=0.042) and WT mice (61.15 ± 1.14% vs. 50.58 ± 2.99 for Test and Train periods, respectively; paired t-test, t=5.597, df=3, p=0.011) had a significant % increase of novel object exploration. This same effect was observed in the discrimination index ratio where values different from zero suggest a preference to explore an object (0.226 ± 0.060 and 0.225 ± 0.023 for Tg2576 and WT, respectively; unpaired t-test, t=0.014, df=7, p=0.989; Supplementary Figure 4C2). We also quantified the total exploration time Tg2576 and WT mice spent exploring the new object (unpaired t-test, t=0.210, df=7, p=0.840; Supplementary Figure 4D1) and the number of approaches the mice had to the novel object (unpaired t-test, t=0.074, df=7, p=0.943; Supplementary Figure 4D2). This lack of significant difference between the two groups suggests that Tg2576 and WT mice had novel object recognition memory.

## DISCUSSION

It is known that the DG contributes to increased excitability in hAPP-overexpressing mouse models of AD [16, 23, 24, 31]. This is important because increased excitability has been implicated in the early stages of AD [30]. Furthermore, the DG has been shown to be a critical regulator of hippocampal hyperexcitability in epilepsy research [47, 51, 90, 91]. Thus, exploring one of the most important cells that regulate GC activity, the MCs, is highly relevant to early stages of AD.

### 1. MCs

Our electrophysiological results suggest that MC excitatory events were greater in Tg2576 mice compared with WT, mainly due to the increased frequency of small events, with no significant changes in mean amplitudes. The Tg2576 cumulative distribution was different from WT, due to the increased number of small events. Interestingly, NMDA receptor-mediated EPSCs appeared to be less affected than AMPA receptor-mediated EPSCs. Tg2576 MC inhibitory events (sIPSCs) were significantly reduced in their mean frequency and amplitude, as well as their cumulative distribution. The data suggest that excitatory input to MCs increases and input from inhibitory neurons decreases in Tg2576 mice. These data are novel because synaptic activity in other studies of mouse models have been reported for older ages and other cell types than MCs [31, 92–95].

It is unclear at the present time what input increases AMPA-EPSCs in MCs and what inhibitory inputs decrease in MCs. Some of the small EPSCs may represent a new input from sprouted axons, since our data suggest the MC axons showed expansion. A new input to MCs might be expected to be small. The depression of inhibitory input could have many sources, but one possibility is that DG interneurons are affected early in life, since they are a major source of inhibitory input to MCs. For instance, alterations in interneurons have been reported to play an important role in AD pathophysiology [31, 92, 94, 95]. Furthermore, there could also be a decline in postsynaptic receptors since sIPSCs decreased in amplitude.

Our results also show changes in intrinsic properties of MCs in Tg2576 mice and most changes that we found would promote excitability. RMP was more depolarized, τ, rheobase and time to the AP peak were significantly reduced, making MCs more prone to fire APs. The lower the time constant values, the faster the membrane will respond to a stimulus, which is consistent with the reduction in the amount of current (rheobase) necessary to trigger APs, and the time it takes for APs to appear after stimulation. However, we also observed a reduction in the peak amplitude of the APs and in the rising slope in Tg2576 MCs, which interestingly, is similar to what has been reported for other cell types in the DG at older ages [95]. The smaller AP amplitudes and slower slopes could be a sign of toxicity due to intracellular Aβ accumulation. Although, another possibility can be that these changes might reflect some compensatory mechanisms to counterbalance the increased MC excitability, as reported for other cell types [95]. Regardless, MC firing behavior showed an enhanced AP firing capacity (F-I curve), as has been shown for other types of hippocampal cells in animal models of AD at older ages [20, 96]. These changes in Tg2576 MCs are consistent with the enhanced E/I ratio for the charge transfer in MCs. All these changes clearly demonstrate enhanced excitability in the Tg2576 mice at 1 month of age.

### 2. GCs

In contrast to what we found for MCs, and in contrast to our previous study of GCs at ∼3 months of age, we did not find any significant difference in the average value of GCs intrinsic properties at 1 month. Thus, WT and Tg2576 GCs were similar in the ability to fire APs, spike frequency adaptation, and intrinsic properties, suggesting that the main cell type, and consequently, the normal function of DG may still not be affected at this very early age. However, spontaneous synaptic activity, in particular the NMDA-mediated excitatory (sEPSCs), and the inhibitory (sIPSCs) inputs to Tg2576 GCs were increased. It is possible that increased excitability of Tg2576 MCs explains the effects because MCs directly excite and indirectly inhibit GCs [64, 69]. Thus, more MC activity would increase excitatory and inhibitory inputs to GCs. However, an increase in MC input to GCs hardly explains why the NMDA-EPSCs of GCs were increased in Tg2576 mice and not the AMPA-EPSCs, since the MC-GC synapses use both AMPA and NMDA receptors. Although, it is known that NMDA receptors are highly concentrated in the IML, in comparison with other parts of the ML [97], where MCs axons are located and innervate GCs [69]. Thus, changes in MC activity can be better reflected in NMDA mediated currents, more than AMPA mediated currents.

### 2. C-Fos protein expression confirms the increased excitability of MCs in Tg2576 mice and little change in Tg2576 GCs

Our electrophysiology data are consistent with the enhanced expression of c-Fos in Tg2576 MCs. MCs seemed to be the main cell type being affected, as a higher number of Tg2576 hilar cells showed enhanced expression of c-Fos, while GCs showed little effect. However, it is relevant to point out the Tg2576 GCL showed higher c-Fos-IF than WT mice even if the number of c-Fos+ GCs were not significantly increased in Tg2576 mice. That result may be due to a greater number of very weakly stained GCs in Tg2576 mice, mostly below the threshold we set for counting bright c-Fos+ cells. This increase in activity markers in the DG has also been reported at older ages in the J20 mouse model of AD, where ΔFosB expression is increased [98]. Furthermore, other cell types in the Tg2576 mice, like septal neurons, have shown enhanced expression of c-Fos after a period of sleep [23], suggesting they are overactive during sleep. The high activity of medial septal neurons is notable because they innervate the DG and could contribute to increased excitability [99–110].

### 3. Role of Aβ

Our data about enhanced intracellular expression of Aβ in 1 month-old Tg2576 hilar cells may explain why MCs increased excitability, as several studies have documented Aβ-induced enhanced excitatory synaptic mechanisms in hippocampal neurons [81, 83, 85, 89, 111]. However, this conclusion should be made with caution because there is also evidence that Aβ increases inhibition [84, 86, 112–114]. Furthermore, we cannot dismiss a contribution of APP and other APP metabolites besides Aβ, as it has been shown they can influence excitability in AD mouse models [115–118].

### 4. MC axon distribution differs between Tg2576 and WT mice at 1 month of age

We and others previously showed that axons from dorsal and ventral MCs differ in their projections in normal conditions [63, 119]. Interestingly, our current study shows that there is a thicker MCs axon bundle in the IML of Tg2576 mice. These changes occurred in sections from a large part of the dorsal-ventral axis, suggesting relatively dorsal and ventral MCs contributed. This small but significant broadening of MC axon bundle may facilitate an increased functional interaction with other inputs to the ML, like the perforant path inputs to GCs, because this appears to occur in normal mice [63, 119]. On the other hand, this expansion of MC axons might also constitute an adaptive mechanism that leads to an enhanced innervation and recruitment of GABAergic interneurons in the ML to compensate for hyperexcitability.

### 5. Tg2576 mice performance in a memory test is intact at 1 month of age

We previously reported the presence of cognitive impairment and alterations in memory performance using novel object tasks in Tg2576 mice at ∼3 months of age [80], which have also been reported in other models of AD at ages older than 3 months [98, 120]. Thus, we wanted to explore whether these physiological and morphological alterations in MCs may be affecting the memory performance at earlier ages in the NOR test, where we know the DG is very relevant. Tg2576 mice did not present any difference compared with WT, suggesting the physiological role of DG is still intact at this age, even when some DG cells (MCs) are already affected. We hypothesize that, ultimately, these changes in the DG will contribute to later impaired cognitive function in Tg2576 mice.

### 6. Sex differences

Sex differences in AD have been of great interest, as AD incidence is greater in female than males [121], and females appear to be affected earlier and more severely in animal models of the disease [122–126]. In the present study, we could not find any significant differences in MCs synaptic and intrinsic properties when sex was considered as factor. However, we observed that the differences in MCs firing activity between genotypes could be seen in males, as females from both genotypes seem to have a larger variability. Regarding GCs, we observed there are some significant differences in their intrinsic properties associated with sex, but no consistent findings that would increase excitability or decrease it. Instead, some sex effects would promote excitability in females and others would reduce it.

## CONCLUSIONS

By using a combination of whole cell electrophysiology and immunohistochemical labeling techniques, this study provided new evidence about the involvement of MCs in the development of the earliest alterations in the Tg2576 mouse model of AD neuropathology. We propose that these changes, involving the enhanced neuronal activity of MCs and the apparent greater innervation of their postsynaptic targets, are among the very first alterations that occur in the DG of Tg2576 mice that, in the long term, lead to the hyperexcitability and the further progression of the pathophysiological process.

## ACKNOWLEDGEMENTS.

We thank John LaFrancois for assistance with mice. Declarations of interest: none.

## ABBREVIATIONS

AD: Alzheimer’s disease.
Aβ: Amyloid β.
DG: Dentate gyrus.
GCs: Granule cells.
MCs: Mossy cells.
MCI: Mild cognitive impairment.
E/I balance: Excitation/Inhibition balance.
sEPSPs: Spontaneous excitatory postsynaptic potentials.
sEPSCs: Spontaneous excitatory postsynaptic currents.
sIPSCs: Spontaneous inhibitory postsynaptic currents.
RMP: Resting membrane potential.
R_in_: Input resistance.
τ: Time constant.
SP: Spike-pair

---

**Supplementary Figure 1. Similar firing behavior in MCs from WT and Tg2576 mice.**

**A.** Spike frequency adaptation was quantified in WT and Tg2576 mice by eliciting a train of 4 APs by positive current steps to MCs. 1. The interspike interval or peak to peak time was measured for each AP pair (spike-pair, SP) and they were plotted sequentially. There was no adaptation and no significant differences by RMANOVA. 2. The first SP and third SP did not show any significant difference between WT and Tg2576 mice.

**B.** Same but trains with 7 APs (6 SPs) were induced. There was no adaptation and there were no significant differences between genotypes. 2. Comparisons of the first and sixth SP showed no significant differences between WT and Tg2576 mice.

**Supplementary Figure 2. GCs firing behavior is not significantly different between WT and Tg2576 mice.**

**A.** The relationship between AP firing frequency and injected current is plotted for WT and Tg2576 mice. Current steps were injected consecutively, and current was increased in 10 pA increments for each step. The mean firing frequency during each step is plotted.

**B.** Spike frequency adaptation was quantified in WT and Tg2576 mice based on a train with 4 APs. 1. The peak to peak time was measured for each AP pair (spike-pair, SP) and plotted sequentially. There were no genotype differences. 2. Statistical comparisons of the first and third SP did not show any significant difference in WT and Tg2576 mice.

**C.** Analogous measurements were made for trains with 7 APs (6 SP). 1. For the sequence of 6 SPs, there were no genotype differences. 2. Comparisons of the first and sixth SP showed significant differences, suggesting GCs from both genotypes adapted their firing, but there were no significant changes between genotypes.

**Supplementary Figure 3. No significant changes in the E/I balance between WT and Tg2576 GCs.**

**A.** The ratio of EPSCs (NMDA- and AMPA-EPSCs) to IPSCs is shown, based on mean frequency (1) or amplitude (2). There were no significant differences between WT and Tg2576 mice.

**B.** Charge transfer of AMPA- and NMDA-sEPSCs (1) and sIPSCs (2) showed no significant difference between genotypes. 3. The E/I ratio for charge transfer based on mean values did not show any differences between Tg2576 and WT mice.

**Supplementary Figure 4. WT and Tg2576 mice showed similar performance on the NOR task at 1 month of age.**

**A.** The experimental timeline for the NOR test. Mice were acclimated (1) for three consecutive days. On the 4^th^ day, mice were subjected to a training session (2) and 1 h later mice were tested in the NOR task (3).

**B.** Quantification of novel object exploration (%) during the training and test session for WT and Tg2576 mice. There was no preference for one object over the other during training, but mice increased exploration of the novel object over the familiar one during testing.

**C.** Quantification of the discrimination index during training (1) and testing (2). There were no genotype differences. During the test, the time mice explored the new object was greater than the familiar object, suggesting a preference to explore the new object.

**D.** 1-2. The time that Tg2576 and WT mice spent exploring the new object (1), and the number of approaches mice had to the object (2) did not show significant differences between genotypes. Tg2576 and WT mice had a similar performance in this task, suggesting that DG function is still intact at this age in the Tg2576 mice.

**Supplementary Table 1. MC intrinsic properties without genotype differences.**

**Supplementary Table 1 legend.**

MC intrinsic properties that showed no significant differences between genotypes are listed. In this table and all others, parametric data were compared using an unpaired Student’s t-test (denoted by the number 1) and non-parametric data were compared by a Mann-Whitney test (denoted by the number 2). Also, the statistical significance was set at p<0.05 and data are presented as mean ± SEM.

**Supplementary Table 2. Sex differences in MC synaptic events.**

**Supplementary Table 2 legend.**

MCs sEPSPs were recorded from WT and Tg2576 male and female mice. A 2-way ANOVA was used to address sex differences, with sex and genotype as factors. There was no effect of sex. There was a significant effect of genotype for sEPSP frequency (boldface font).

**Supplementary table 3. Sex differences in MC intrinsic properties.**

**Supplementary table 3 legend.**

Values of the different intrinsic properties obtained from WT and Tg2576 MCs from male and female mice. Statistical comparisons for evaluating sex differences were performed using a 2- way ANOVA with sex and genotype as factors. There were no significant effects of sex but there were effects of genotype (boldface font).

**Supplementary Table 4. GC synaptic events without genotype differences.**

**Supplementary Table 4 legend.**

The results of GC recordings of sEPSPs and AMPA-sEPSCs are shown. There were no significant differences between WT and Tg2576 mice.

**Supplementary Table 5. GC intrinsic properties without genotype differences.**

**Supplementary Table 5 legend.**

The results of GC recordings of intrinsic properties are shown. There were no significant differences between WT and Tg2576 mice.

**Supplementary Table 6. Sex differences in GC synaptic events.**

**Supplementary Table 6 legend.**

Results of GC recordings of synaptic events (sEPSPs, AMPA-sEPSCs, NMDA-sEPSCs and sIPSCs) from WT and Tg2576 mice. A 2-way ANOVA was used to evaluate sex differences with sex and genotype as factors. There were significant effects of sex on AMPA-sEPSC amplitude and sIPSC frequency. For AMPA-sEPSC amplitude, there was an interaction of factors with the amplitudes higher for WT females than WT males but lower for Tg2576 females compared to Tg2576 males. For sIPSC frequency, there was greater frequency in female Tg2576 females compared to Tg2576 males. However, WT males and females were not significantly different.

**Supplementary Table 7. Sex differences in GC intrinsic properties.**

**Supplementary Table 7 legend.**

Results of GC recordings of intrinsic properties from WT and Tg2576 mice. A 2-way ANOVA was used to evaluate sex differences with sex and genotype as factors. There was a significant effect of sex on R_in_, rheobase, half-width, and the maximum slope of the AP decay phase. There was an interaction of factors for threshold and half-width. For R_in_, females had lower values. For rheobase, females had higher values. For half-width, females had shorter half-widths than males but only in Tg2576 mice. For the slope of the AP decay, females had a faster repolarization in both genotypes. For threshold, female WT mice had a more hyperpolarized threshold compared to male WT mice, and female Tg2576 mice had a more depolarized threshold than male Tg2576 mice. Thus, there were diverse sex differences, but excitability was not consistently higher in one sex.

