## Supplementary Figures and Tables for "Increased excitability of dentate gyrus mossy cells occurs early in life in the Tg2576 model of Alzheimer’s disease"

### Supplementary Figure 1. MC firing behavior.

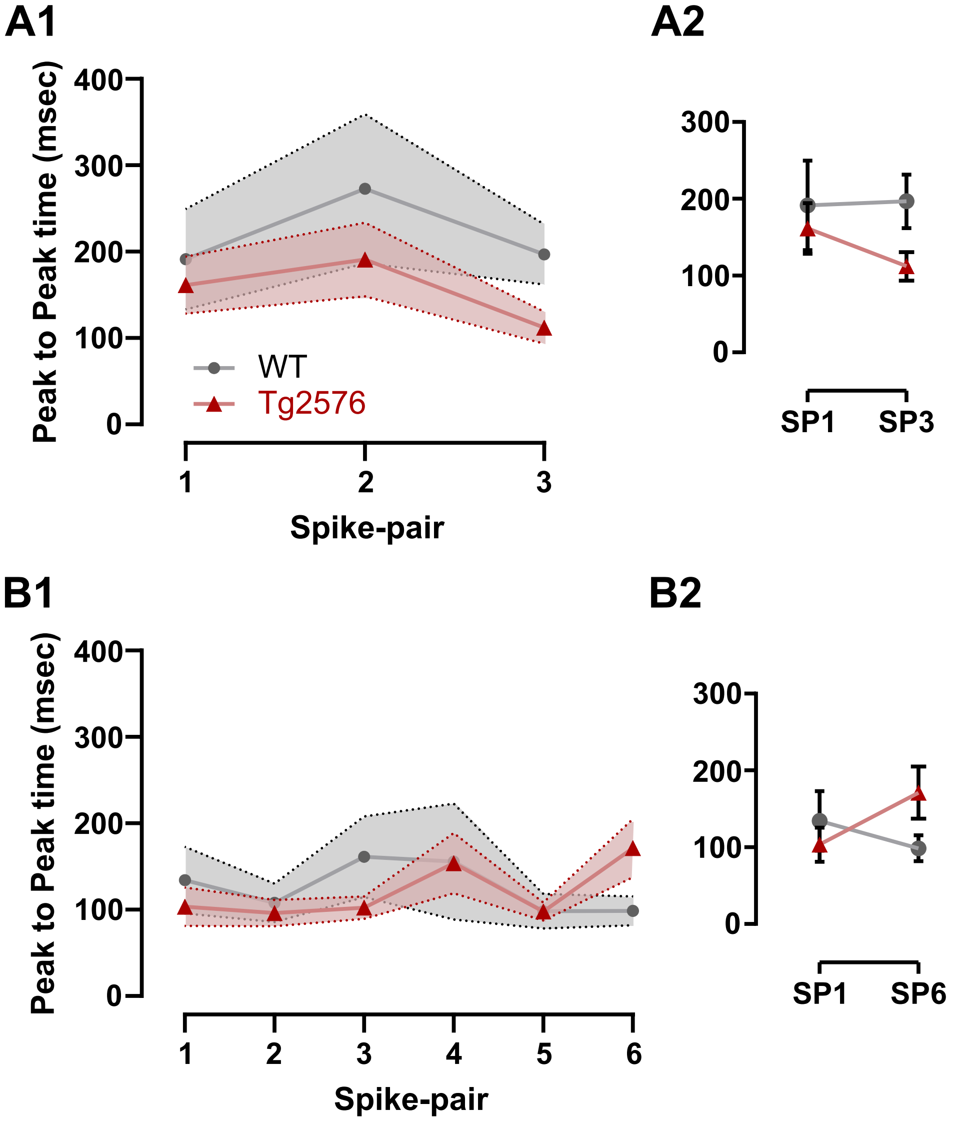

### Supplementary Figure 2. GC firing behavior.

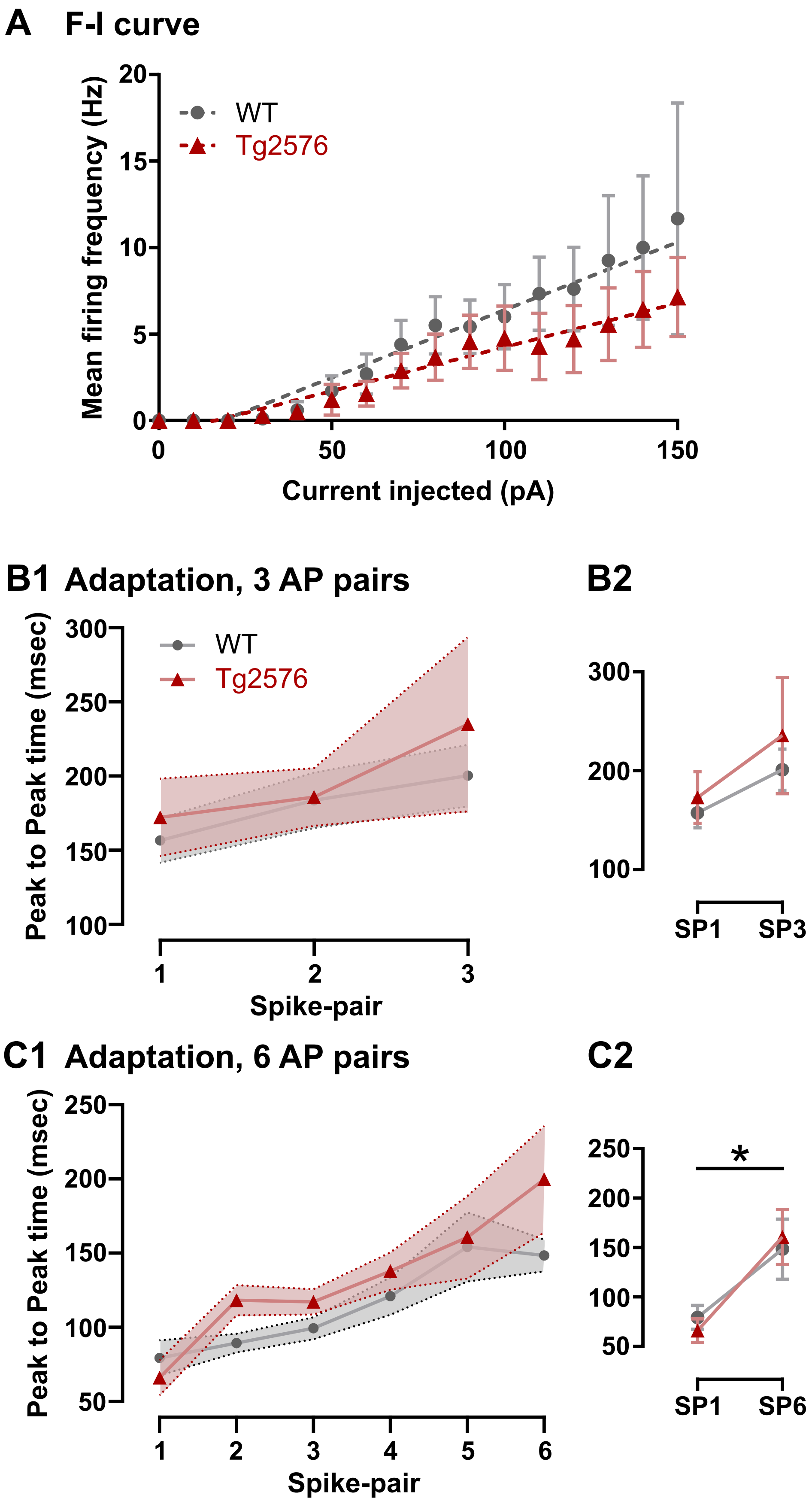

### Supplementary Figure 3. GC E/I balance.

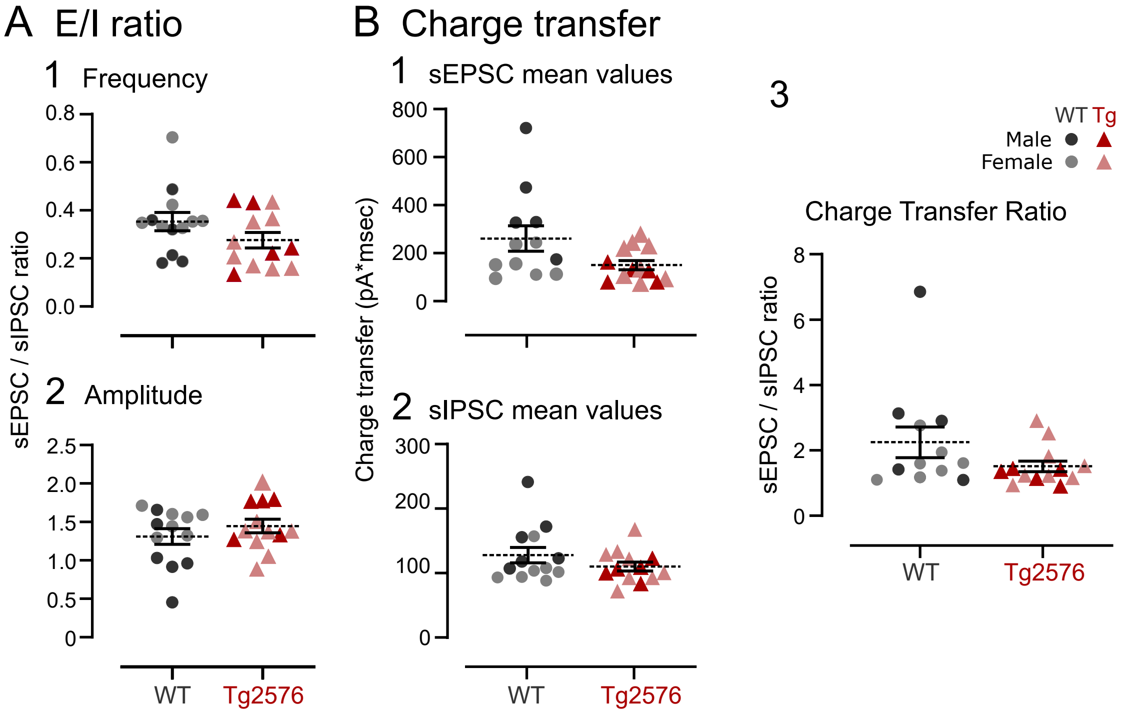

### Supplementary Figure 4. NOR at 1 month of age.

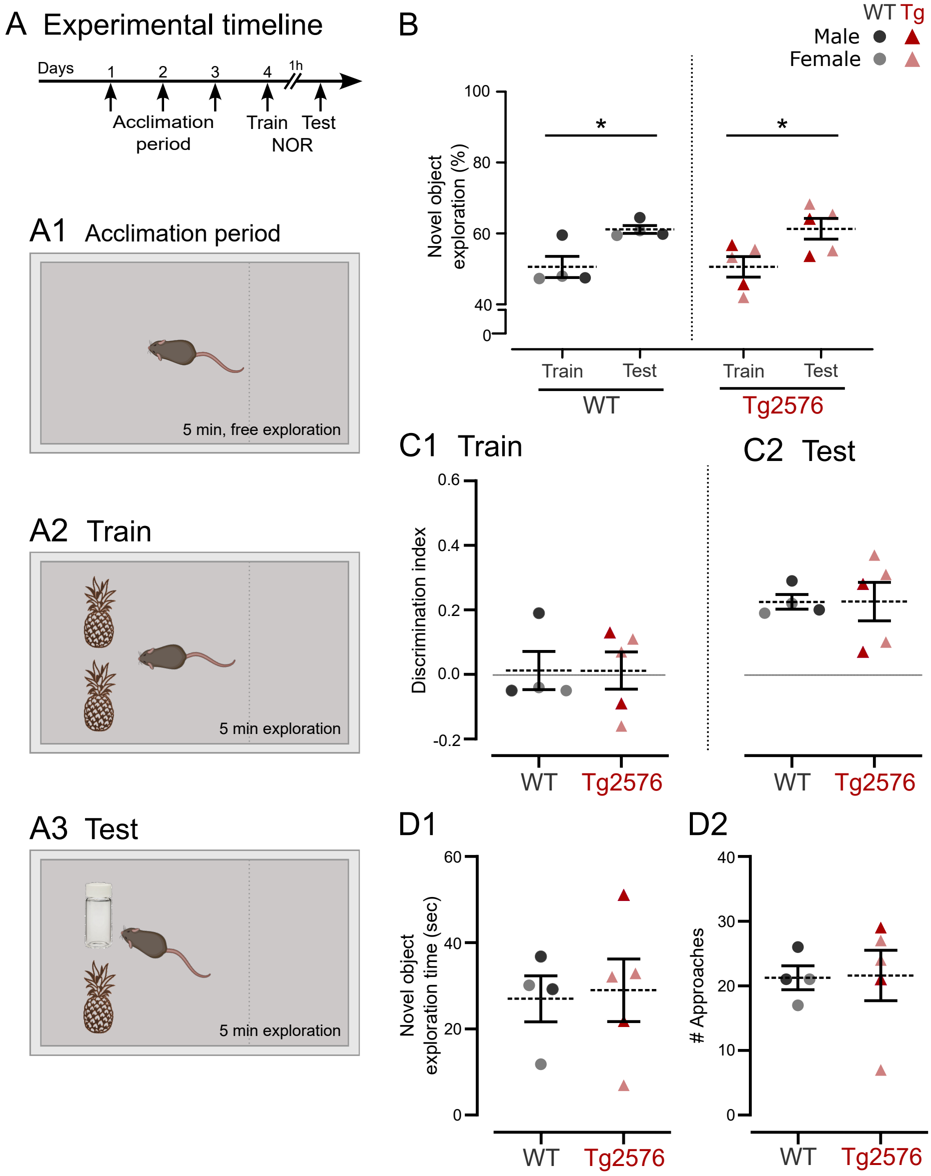

**Supplementary Table 1. MC intrinsic properties without genotype differences.**

| | $R_{in}$ (MΩ) | Threshold (mV) | Time to peak from threshold (msec) |
|---|---|---|---|
| **WT** | 320.50 ± 40.64 | -43.92 ± 2.55 | 0.88 ± 0.08 |
| **Tg2576** | 279.30 ± 35.52 | -47.35 ± 2.74 | 1.01 ± 0.15 |
| **P value** | 0.451 | 0.370 | 0.515 |
| **Statistical Test** | U=87 (2) | t=0.916, df=21 (1) | U=55 (2) |
| **$N_{WT}$** | 15 | 12 | 12 |
| **$N_{Tg2576}$** | 14 | 11 | 11 |

| | Half-width (msec) | Max Decay Slope | dv/dt ratio |
|---|---|---|---|
| **WT** | 0.42 ± 0.05 | -62.05 ± 4.15 | -2.64 ± 0.27 |
| **Tg2576** | 0.43 ± 0.04 | -51.88 ± 5.75 | -2.16 ± 0.05 |
| **P value** | 0.661 | 0.161 | 0.379 |
| **Statistical Test** | U=58.5 (2) | t=1.453, df=21 (1) | U=51 (2) |
| **$N_{WT}$** | 12 | 12 | 12 |
| **$N_{Tg2576}$** | 11 | 11 | 11 |

**(1) Parametric test. Unpaired t-test.**

**(2) Nonparametric test. Mann-Whitney test.**

**Supplementary Table 2. Sex differences in MC synaptic events.**

| | Frequency | | Amplitude | |
|---|---|---|---|---|
| **sEPSPs** | **WT** | **Tg2576** | **WT** | **Tg2576** |
| **Male** | 5.86 ± 0.66 | 8.42 ± 1.45 | 2.73 ± 0.55 | 2.26 ± 0.33 |
| **Female** | 5.81 ± 0.61 | 9.23 ± 1.25 | 2.21 ± 0.44 | 1.88 ± 0.22 |
| **$N_{male}$** | 11 | 9 | 11 | 9 |
| **$N_{female}$** | 7 | 9 | 7 | 9 |
| **2-way ANOVA (P value)** | Interaction | 0.693 | Interaction | 0.882 |
| | Sex | 0.727 | Sex | 0.312 |
| | Genotype | **0.009** | Genotype | 0.369 |
| | | **$F_{(1, 32)} = 7.698$** | | |

**Supplementary Table 3. Sex differences in MC intrinsic properties.**

| | RMP | | Tau | |
|---|---|---|---|---|
| | WT | Tg2576 | WT | Tg2576 |
| **Male** | -72.13 ± 1.91 | -58.63 ± 3.31 | 49.76 ± 3.77 | 42.15 ± 5.63 |
| **Female** | -70.61 ± 0.91 | -60.90 ± 1.44 | 54.94 ± 10.41 | 37.08 ± 4.87 |
| **N male** | 11 | 9 | 11 | 7 |
| **N female** | 5 | 8 | 4 | 6 |
| | Interaction | 0.435 | Interaction | 0.383 |
| **2-way ANOVA (P value)** | Sex | 0.877 | Sex | 0.993 |
| | Genotype | **<0.001** | Genotype | **0.037** |
| | | **F (1, 29) = 23.50** | | **F (1, 24) = 4.880** |

| | R in | | Rheobase | |
|---|---|---|---|---|
| | WT | Tg2576 | WT | Tg2576 |
| **Male** | 333.30 ± 51.13 | 303.0 ± 49.58 | 100.00 ± 14.88 | 44.29 ± 5.28 |
| **Female** | 285.50 ± 66.24 | 247.6 ± 52.10 | 147.50 ± 64.47 | 61.00 ± 17.20 |
| **N male** | 11 | 8 | 8 | 7 |
| **N female** | 4 | 6 | 4 | 5 |
| | Interaction | 0.950 | Interaction | 0.544 |
| **2-way ANOVA (P value)** | Sex | 0.340 | Sex | 0.213 |
| | Genotype | 0.572 | Genotype | **0.010** |
| | | | | **F (1, 20) = 8.123** |

| | Threshold | | Peak Amplitude | |
|---|---|---|---|---|
| | WT | Tg2576 | WT | Tg2576 |
| **Male** | -46.81 ± 3.24 | -49.38 ± 3.68 | 91.49 ± 4.52 | 77.18 ± 5.49 |
| **Female** | -38.16 ± 2.41 | -43.91 ± 2.99 | 97.52 ± 4.61 | 79.37 ± 5.69 |
| **N male** | 8 | 7 | 8 | 7 |
| **N female** | 4 | 5 | 4 | 5 |
| | Interaction | 0.659 | Interaction | 0.728 |
| **2-way ANOVA (P value)** | Sex | 0.061 | Sex | 0.460 |
| | Genotype | 0.255 | Genotype | **0.008** |
| | | | | **F (1, 20) = 8.859** |

| | Time to Peak | | Time to Peak from threshold | |
|---|---|---|---|---|
| | WT | Tg2576 | WT | Tg2576 |
| **Male** | 582.60 ± 68.02 | 306.9 ± 78.66 | 0.88 ± 0.10 | 1.01 ± 0.07 |
| **Female** | 429.30 ± 66.65 | 349.0 ± 83.73 | 0.90 ± 0.11 | 1.10 ± 0.40 |
| **N male** | 8 | 7 | 8 | 7 |
| **N female** | 4 | 4 | 4 | 4 |
| | Interaction | 0.245 | Interaction | 0.869 |
| **2-way ANOVA (P value)** | Sex | 0.503 | Sex | 0.758 |
| | Genotype | **0.042** | Genotype | 0.342 |
| | | **F (1, 19) = 4.778** | | |

| | Half-width | | Max Rise Slope | |
|---|---|---|---|---|
| | WT | Tg2576 | WT | Tg2576 |
| **Male** | 0.37 ± 0.04 | 0.41 ± 0.02 | 154.70 ± 20.73 | 103.60 ± 12.15 |
| **Female** | 0.51 ± 0.12 | 0.47 ± 0.11 | 180.10 ± 36.02 | 129.50 ± 35.10 |
| **N male** | 8 | 7 | 8 | 7 |
| **N female** | 4 | 4 | 4 | 4 |
| | Interaction | 0.565 | Interaction | 0.991 |
| **2-way ANOVA (P value)** | Sex | 0.139 | Sex | 0.313 |
| | Genotype | 0.999 | Genotype | 0.054 |

| | Max Decay Slope | | dv/dt ratio | |
|---|---|---|---|---|
| | WT | Tg2576 | WT | Tg2576 |
| **Male** | -64.54 ± 5.71 | -48.57 ± 5.65 | -2.35 ± 0.18 | -2.13 ± 0.03 |
| **Female** | -57.07 ± 4.99 | -57.68 ± 13.17 | -3.23 ± 0.68 | -2.21 ± 0.15 |
| **N male** | 8 | 7 | 8 | 7 |
| **N female** | 4 | 4 | 4 | 4 |
| | Interaction | 0.282 | Interaction | 0.167 |
| **2-way ANOVA (P value)** | Sex | 0.914 | Sex | 0.101 |
| | Genotype | 0.318 | Genotype | **0.038** |
| | | | | **F (1, 19) = 4.960** |

**Supplementary Table 4. GC synaptic events without genotype differences.**

| | sEPSPs frequency (events/sec) | sEPSPs amplitude (mV) |
|---|---|---|
| **WT** | 1.73 ± 0.15 | 0.46 ± 0.04 |
| **Tg2576** | 2.06 ± 0.24 | 0.46 ± 0.04 |
| **P value** | 0.487 | 0.996 |
| **Statistical Test** | U=54 (2) | t=0.006, df=21 (1) |
| **N $_{WT}$** | 11 | 11 |
| **N $_{Tg2576}$** | 12 | 12 |
| | **AMPA-sEPSCs frequency (events/sec)** | **AMPA-sEPSCs amplitude (pA)** |
| **WT** | 1.78 ± 0.17 | 7.21 ± 0.71 |
| **Tg2576** | 2.00 ± 0.30 | 8.02 ± 0.85 |
| **P value** | 0.534 | 0.471 |
| **Statistical Test** | t=0.631, df=24 (1) | t=0.733, df=24 (1) |
| **N $_{WT}$** | 13 | 13 |
| **N $_{Tg2576}$** | 13 | 13 |

**(1) Parametric test. Unpaired t-test.**
**(2) Nonparametric test. Mann-Whitney test.**

**Supplementary Table 5. GC intrinsic properties without genotype differences.**

| | RMP (mV) | Tau (msec) | $R_{in}$ (MΩ) | Rheobase (mV) |
|---|---|---|---|---|
| WT | -78.85 ± 2.66 | 33.92 ± 2.46 | 550.40 ± 43.72 | 65.00 ± 8.06 |
| Tg2576 | -76.11 ± 2.95 | 30.74 ± 4.32 | 405.80 ± 64.17 | 84.00 ± 14.85 |
| P value | 0.499 | 0.531 | 0.079 | 0.276 |
| Statistical Test | t=0.688, df=18 (1) | t=0.639, df=18 (1) | t=1.862, df=18 (1) | t=1.125, df=18 (1) |
| $N_{WT}$ | 10 | 10 | 10 | 10 |
| $N_{Tg2576}$ | 10 | 10 | 10 | 10 |

| | Threshold (mV) | Peak Amplitude (mV) | Time to Peak (msec) | Half-width (msec) |
|---|---|---|---|---|
| WT | -36.90 ± 2.62 | 102.20 ± 3.96 | 506.20 ± 91.31 | 0.38 ± 0.02 |
| Tg2576 | -36.24 ± 2.04 | 109.80 ± 3.66 | 529.40 ± 88.61 | 0.49 ± 0.05 |
| P value | 0.436 | 0.179 | 0.858 | 0.052 |
| Statistical Test | U = 39 (2) | t=1.399, df=18 (1) | t=0.182, df=18 (1) | t=2.083, df=18 (2) |
| $N_{WT}$ | 10 | 10 | 10 | 10 |
| $N_{Tg2576}$ | 10 | 10 | 10 | 10 |

| | Max Rise Slope (mV/msec) | Max Decay Slope (mV/msec) | dv/dt ratio |
|---|---|---|---|
| WT | 137.10 ± 12.03 | -62.56 ± 5.18 | 2.23 ± 0.15 |
| Tg2576 | 133.50 ± 11.49 | -53.16 ± 3.81 | 2.56 ± 0.21 |
| P value | 0.831 | 0.161 | 0.190 |
| Statistical Test | t=0.2165, df=18 (1) | t=1.462, df=18 (1) | U = 32 (2) |
| $N_{WT}$ | 10 | 10 | 10 |
| $N_{Tg2576}$ | 10 | 10 | 10 |

**(1) Parametric test. Unpaired t-test.**
**(2) Nonparametric test. Mann-Whitney test.**

**Supplementary Table 6. Sex differences in GC synaptic events.**

| | sEPSPs frequency | | sEPSPs amplitude | |
|---|---|---|---|---|
| | WT | Tg2576 | WT | Tg2576 |
| **Male** | 1.84 ± 0.20 | 1.71 ± 0.33 | 0.46 ± 0.06 | 0.46 ± 0.08 |
| **Female** | 1.44 ± 0.10 | 2.31 ± 0.31 | 0.48 ± 0.05 | 0.464 ± 0.04 |
| **N male** | 8 | 5 | 8 | 5 |
| **N female** | 3 | 7 | 3 | 7 |
| **2-way ANOVA (P value)** | Interaction | 0.111 | Interaction | 0.874 |
| | Sex | 0.744 | Sex | 0.868 |
| | Genotype | 0.234 | Genotype | 0.945 |

| | AMPA-sEPSCs frequency | | AMPA-sEPSCs amplitude | |
|---|---|---|---|---|
| | WT | Tg2576 | WT | Tg2576 |
| **Male** | 1.56 ± 0.34 | 1.50 ± 0.38 | 4.88 ± 0.45 | 8.15 ± 1.93 |
| **Female** | 1.97 ± 0.12 | 2.31 ± 0.41 | 9.21 ± 0.55 | 7.94 ± 0.82 |
| **N male** | 6 | 5 | 6 | 5 |
| **N female** | 7 | 8 | 7 | 8 |
| **2-way ANOVA (P value)** | Interaction | 0.572 | **Interaction** | **0.029**<br>**$F_{(1, 22)} = 5.426$** |
| | Sex | 0.086 | **Sex** | **0.045**<br>**$F_{(1, 22)} = 4.506$** |
| | Genotype | 0.681 | Genotype | 0.314 |

| | NMDA-sEPSCs frequency | | NMDA-sEPSCs amplitude | |
|---|---|---|---|---|
| | WT | Tg2576 | WT | Tg2576 |
| **Male** | 0.83 ± 0.18 | 1.13 ± 0.16 | 7.32 ± 0.61 | 7.35 ± 1.30 |
| **Female** | 0.95 ± 0.09 | 1.56 ± 0.26 | 8.99 ± 0.72 | 8.97 ± 1.06 |
| **N male** | 5 | 5 | 5 | 5 |
| **N female** | 7 | 8 | 7 | 8 |
| **2-way ANOVA (P value)** | Interaction | 0.459 | Interaction | 0.985 |
| | Sex | 0.195 | Sex | 0.116 |
| | Genotype | **0.035**<br>**$F_{(1, 21)} = 5.076$** | Genotype | 0.995 |

| | sIPSCs frequency | | sIPSCs amplitude | |
|---|---|---|---|---|
| | WT | Tg2576 | WT | Tg2576 |
| **Male** | 7.37 ± 0.84 | 9.28 ± 0.85 | 10.57 ± 1.58 | 9.42 ± 1.38 |
| **Female** | 7.63 ± 0.67 | 15.27 ± 1.95 | 12.09 ± 0.55 | 12.87 ± 1.41 |
| **N male** | 6 | 5 | 6 | 5 |
| **N female** | 7 | 8 | 7 | 8 |
| **2-way ANOVA (P value)** | Interaction | 0.051 | Interaction | 0.469 |
| | **Sex** | **0.034**<br>**$F_{(1, 22)} = 5.097$** | Sex | 0.071 |
| | **Genotype** | **0.002**<br>**$F_{(1, 22)} = 11.89$** | Genotype | 0.889 |

**Supplementary Table 7. Sex differences in GC intrinsic properties.**

| | RMP | | tau | |
|---|---|---|---|---|
| | WT | Tg2576 | WT | Tg2576 |
| **Male** | -77.42 ± 3.20 | -76.80 ± 4.33 | 34.00 ± 2.79 | 38.20 ± 7.18 |
| **Female** | -82.18 ± 5.15 | -75.43 ± 4.49 | 33.75 ± 6.00 | 23.29 ± 2.17 |
| **N male** | 7 | 5 | 7 | 5 |
| **N female** | 3 | 5 | 3 | 5 |
| **2-way ANOVA (P value)** | Interaction | 0.489 | Interaction | 0.145 |
| | Sex | 0.700 | Sex | 0.132 |
| | Genotype | 0.407 | Genotype | 0.522 |

| | Rin | | Rheobase | |
|---|---|---|---|---|
| | WT | Tg2576 | WT | Tg2576 |
| **Male** | 591.60 ± 50.04 | 518.60 ± 96.32 | 57.14 ± 6.80 | 56.00 ± 14.70 |
| **Female** | 454.30 ± 67.34 | 293.00 ± 53.72 | 83.33 ± 20.28 | 112.0 ± 19.60 |
| **N male** | 7 | 5 | 7 | 5 |
| **N female** | 3 | 5 | 3 | 5 |
| **2-way ANOVA (P value)** | Interaction | 0.546 | Interaction | 0.335 |
| | Sex | **0.022** **$F_{(1, 16)} = 6.423$** | Sex | **0.015** **$F_{(1, 16)} = 7.515$** |
| | Genotype | 0.121 | Genotype | 0.372 |

| | Threshold | | Peak Amplitude | |
|---|---|---|---|---|
| | WT | Tg2576 | WT | Tg2576 |
| **Male** | -34.38 ± 3.35 | -40.76 ± 2.43 | 100.60 ± 4.99 | 109.80 ± 6.74 |
| **Female** | -42.76 ± 0.54 | -31.72 ± 1.62 | 106.10 ± 7.08 | 109.70 ± 3.84 |
| **N male** | 7 | 5 | 7 | 5 |
| **N female** | 3 | 5 | 3 | 5 |
| **2-way ANOVA (P value)** | Interaction | **0.010** **$F_{(1, 16)} = 8.624$** | Interaction | 0.642 |
| | Sex | 0.913 | Sex | 0.654 |
| | Genotype | 0.444 | Genotype | 0.295 |

| | Time to peak | | Half-width | |
|---|---|---|---|---|
| | WT | Tg2576 | WT | Tg2576 |
| **Male** | 513.00 ± 83.05 | 601.3 ± 132.7 | 0.39 ± 0.03 | 0.61 ± 0.03 |
| **Female** | 490.50 ± 276.8 | 457.4 ± 123.0 | 0.37 ± 0.05 | 0.37 ± 0.03 |
| **N male** | 7 | 7 | 7 | 5 |
| **N female** | 3 | 3 | 3 | 5 |
| **2-way ANOVA (P value)** | Interaction | 0.667 | Interaction | **0.005** **$F_{(1, 16)} = 10.92$** |
| | Sex | 0.557 | Sex | **0.002** **$F_{(1, 16)} = 13.67$** |
| | Genotype | 0.845 | Genotype | **0.006** **$F_{(1, 16)} = 9.949$** |

| | Max Rise Slope | | Max Decay Slope | |
|---|---|---|---|---|
| | WT | Tg2576 | WT | Tg2576 |
| **Male** | 124.80 ± 13.29 | 121.1 ± 9.82 | -55.02 ± 5.00 | -49.44 ± 6.31 |
| **Female** | 166.00 ± 18.47 | 146.0 ± 20.49 | -80.16 ± 2.50 | -56.88 ± 4.30 |
| **N male** | 7 | 5 | 7 | 5 |
| **N female** | 3 | 5 | 3 | 5 |
| **2-way ANOVA (P value)** | Interaction | 0.627 | Interaction | 0.131 |
| | Sex | 0.062 | Sex | **0.010** **$F_{(1, 16)} = 8.605$** |
| | Genotype | 0.482 | Genotype | **0.019** **$F_{(1, 16)} = 6.75$** |

| | dv/dt ratio | |
|---|---|---|
| | WT | Tg2576 |
| **Male** | -2.30 ± 0.20 | -2.60 ± 0.31 |
| **Female** | -2.06 ± 0.16 | -2.52 ± 0.23 |
| **N male** | 7 | 5 |
| **N female** | 3 | 5 |
| **2-way ANOVA (P value)** | Interaction | 0.774 |
| | Sex | 0.580 |
| | Genotype | 0.201 |
